# Plasma cell maintenance and antibody secretion are under the control of Sec22b-mediated regulation of organelle dynamics

**DOI:** 10.1101/2022.01.14.476154

**Authors:** Amélie Bonaud, Laetitia Gargowitsch, Simon M. Gilbert, Elanchezhian Rajan, Pablo Canales Herrerias, Daniel Stockholm, Nabila F. Rahman, Mark O. Collins, Danika L. Hill, Andres Alloatti, Nagham Alouche, Stéphanie Balor, Vanessa Soldan, Daniel Gillet, Julien Barbier, Françoise Bachelerie, Kenneth G.C. Smith, Pierre Bruhns, Sebastian Amigorena, Karl Balabanian, Michelle A. Linterman, Andrew A. Peden, Marion Espéli

**Affiliations:** Université de Paris, Institut de Recherche Saint-Louis, INSERM U1160, F-75010 Paris, France; CNRS, GDR3697 “Microenvironment of tumor niches”, Micronit, France; OPALE Carnot Institute, The Organization for Partnerships in Leukemia, Hôpital Saint-Louis, Paris, France; Université Paris-Saclay, INSERM, Inflammation, Microbiome and Immunosurveillance, Clamart, France; Department of Medicine, University of Cambridge, Cambridge Biomedical Campus, Addenbrooke’s Hospital, Cambridge, UK; School of Bioscience, University of Sheffield, Western Bank, Sheffield, S102TN, UK; Institut Pasteur, Université de Paris, Unit of Antibodies in Therapy and Pathology, Inserm UMR1222, F-75015 Paris; PSL Research University, EPHE, Paris, France; Sorbonne Université, INSERM, Centre de Recherche Saint-Antoine, CRSA, F-75012, Paris, France; Dementia Research Institute, University of Cardiff, Hadyn Ellis Building, Maindy Road, Cardiff, CF24 4HQ; Lymphocyte Signalling and Development, Babraham Institute, Babraham Research Campus, Cambridge CB22 3AT, UK; Department of Immunology and Pathology, Monash University, Melbourne, Victoria, 3004, Australia; PSL Research University, Institut Curie Research Center, INSERM U932, Paris, France; Facultad de Ciencias Médicas, Instituto de Inmunología Clínica y Experimental de Rosario (IDICER)-CONICET/Universidad Nacional de Rosario, Rosario, Argentina; METi, Centre de Biologie Intégrative, Université de Toulouse, CNRS, UPS, 31062, Toulouse, France; Université Paris-Saclay, CEA, INRAE, Département Médicaments et Technologies pour la Santé (DMTS), SIMoS, 91191 Gif-sur-Yvette, France; Jeffrey Cheah Biomedical Centre Cambridge Biomedical, Cambridge Institute of Therapeutic Immunology & Infectious Disease, University of Cambridge, Cambridge, United Kingdom

**Keywords:** Plasma cell, antibody, SNARE - Endoplasmic reticulum, Mitochondria

## Abstract

Despite the essential role of plasma cells in health and disease, the cellular mechanisms controlling their survival and secretory capacity are still poorly understood. Here, we identified the SNARE Sec22b as a unique and critical regulator of plasma cell maintenance and function. In absence of Sec22b, plasma cells were barely detectable and serum antibody titres were dramatically reduced. Accordingly, *Sec22b* deficient mice fail to mount a protective immune response. At the mechanistic level, we demonstrated that Sec22b is indispensable for efficient antibody secretion but also for plasma cell fitness through the regulation of the morphology of the endoplasmic reticulum and mitochondria. Altogether, our results unveil a critical role for Sec22b-mediated regulation of plasma cell biology through the control of organelle dynamics.

## Introduction

Plasma cells (PCs) are the cellular source of humoral immunity via the long-term secretion of large quantities of antibodies that provide protection against reinfection. These cells can also contribute to diseases including plasmacytomas as well as antibody-mediated autoimmune and inflammatory pathologies. However, the therapeutic arsenal to target PCs is still very limited. Despite the essential role of PCs in health and disease, the cellular mechanisms controlling their secretory function and their survival are poorly understood. Closing this knowledge gap is thus of paramount importance for designing new approaches to target this cell type.

During the transition from B cell to PC, the cell is reprogrammed to produce and secrete around 10^2^-10^3^ antibodies per second^1^. To accommodate this large protein load, PCs expand their endoplasmic reticulum (ER) and adapt to tolerate the extra stress induced via upregulation of the Ire1α/Xbp1 branch of the unfolded protein response (UPR)^2–4^. Early works from the 70s report that antibody secretion happens via the conventional constitutive exocytosis pathway meaning that they are not pre-stocked in granules but secreted as they are produced through the Golgi apparatus where they are glycosylated^5–7^. Antibody transport from the expanded ER to the Golgi apparatus is thus the key bottleneck in this process, however, the molecular mechanisms at play are still unknown^8^.

Members of the Soluble N-ethylmaleimide-sensitive factor Attachment Protein Receptor (SNARE) family are essential for the intracellular transport and fusion of protein cargoes between organelles. They are involved in both regulated and constitutive exocytosis and form a large family composed of different subtypes (Qa-, Qb-, Qc- and R-SNAREs)^9^. Distinct sets of SNAREs are expressed on the different organelles and can also be cell type specific^10^. On top of vesicular transport, SNAREs can have other non-canonical functions, some of which could be highly relevant for PC biology, including ER branching, regulation of autophagosome maturation as well as plasma membrane expansion^11–16^.

In this work we explore the role of SNARE proteins required for ER to Golgi transport, and in particular of Sec22b, in antibody secretion and PC biology. In absence of Sec22b PCs are dramatically reduced. The transport of antibodies from the ER to the Golgi is impacted leading to a strong decrease in antibody secretion and an incapacity to generate a protective humoral response. This defect in antibody production and PC survival is accompanied by deregulation of the UPR and a defective ER morphology.

In addition, we observed that in absence of Sec22b the mitochondrial network is hyperfused suggesting a key role for this protein in the regulation of ER-mediated mitochondrial fission. Altogether our results establish Sec22b as a critical regulator of PC biology through the control of conventional constitutive secretion but also through the regulation of organelle morphology and dynamics.

## Results

### The Syntaxin-5/Sec22b SNARE complex is overexpressed in plasma cells

To identify new molecular actors that contribute to antibody secretion and PC fitness we analysed the expression of molecules involved in ER to Golgi transport by mining a proteogenomic database using the Plasmacytomics interface (https://plasmacytomics.shinyapps.io/home/; manuscript in preparation). To determine the validity of the platform we first confirmed that the expression of well-known factors of B cell and PC differentiation behaved as one would predict. As expected, the B cell transcription factor Pax5 was significantly downregulated in PCs compared to B cells while genes encoding the transcription factor Xbp1 and the PC marker CD138 (*Sdc1*) were significantly upregulated in PCs (Figure 1A). Molecules involved in ER to Golgi trafficking, including SNAREs (Syntaxin 5, Sec22B and Ykt6), a tethering factor (USO1/p115) and the Sec1/Munc-18 protein (SCFD1/Sly1), were upregulated at the transcriptional level in PCs compared to B cells (Figure 1A). In support of the transcriptomic data, we also observed a significant increase in the levels of many of these proteins by mass-spectrometry (Figure 1B and Supplementary Figure 1) and immunoblotting (Figure 1C). Datamining of human RNAseq datasets using the plasmacytomics platform also showed significantly increased expression of *SEC22B* and *STX5A* in human PCs compared to B cells (Figure 1D) suggesting that these changes are conserved across species.

**Figure 1:**
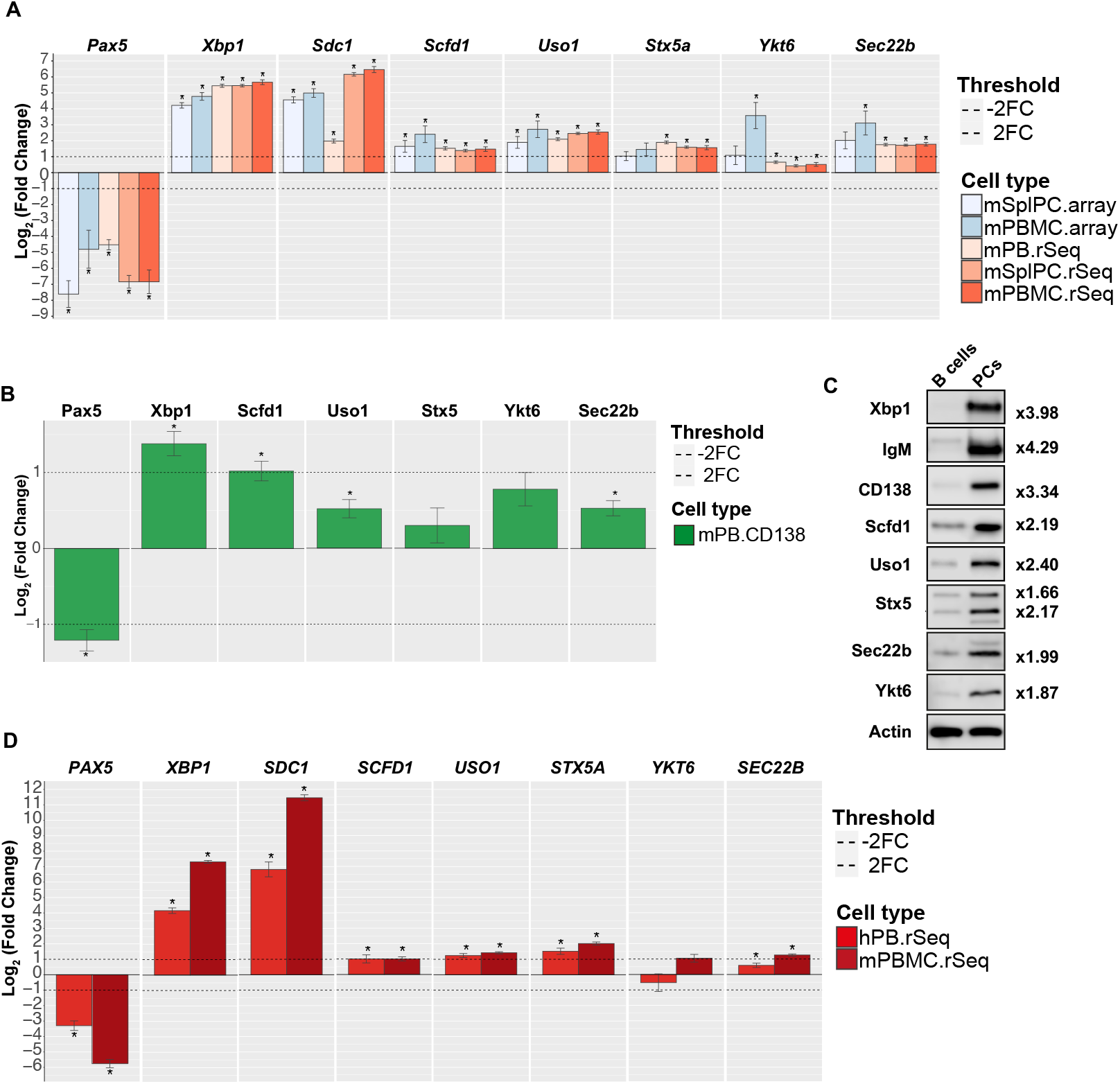
The Syntaxin-5-Sec22b SNARE complex is overexpressed in plasma cells: A) The PlasmacytOMICs interface (manuscript under preparation) was used to perform a meta-analysis of gene expression changes between murine Naive B cells and a range of antibody secreting cell types (mSplPC.Array = microarray / mouse splenic PCs; mBMPC.Array = microarray / mouse BM PCs; mPB.rSeq = RNAseq / mouse plasmablasts generated *in vitro*; mSplPC.rSeq = RNAseq / mouse splenic PCs; mBMPC.rSeq = RNAseq / mouse BM PCs) for the indicated genes. The Log_2_ Fold change between naïve B cells and the indicated cell subset is shown for each gene. Error bars show SEM. * - indicates False Discovery Adjusted p-value < 0.05. B) Changes in protein abundance between murine Naive B cells and CD138 enriched PCs generated in vitro were measured using label free LC-MS/MS analysis and plotted using the PlasmacytOMICcs interface. The Log_2_ Fold change between naïve B cells and the indicated cell subset is shown for each protein. Error bars show SEM. * - indicates False Discovery Adjusted p-value < 0.05. We were unable to consistently measure sufficient peptides for quantification of CD138 in the PC samples possibly due to its high level of glycosylation. The fold change shown for XBP1 is an underestimate, as the ratio plotted is calculated using an imputed value for the B-cell samples as the protein was not detected in these samples (see Supplementary Figure 1). C) Representative immunoblots for Xbp1, IgM, CD138, Scfd1, Uso1, Stx5, Ytk6, Sec22b and b-actin (top to bottom respectively) from samples prepared from splenic B cells (left panel) or CD138 enriched *in vitro* differentiated PCs (right panel). The band shown for CD138 is the non-glycosylated form of the protein. For Stx5 the two bands correspond to the short and the long isoforms of the protein. The fold change between B cells and PCs normalized to b-actin is indicated on the right for each protein. D) Changes in gene expression between human Naive B cells and selection of human antibody secreting cell types (hPB.rSeq = RNAseq / human blood PC; hBMPC.rSeq = RNAseq / human BM PC) were calculated for the indicated genes using the PlasmacytOMICcs platform. The Log_2_ Fold change between naïve B cells and the indicated cell subset is shown for each gene. Error bars show SEM. * - indicates False Discovery Adjusted p-value < 0.05.

Previous works has shown that the Golgi-localised SNARE, Stx5 forms a complex with the ER-localised Sec22b to allow fusion of transport vesicles between these two organelles^10,17,18^. To test the functional relevance of Stx5 expression in PC biology, we developed a cell culture and differentiation assay using control shRNA or an shRNA specific for Stx5 together with GFP reporter expression (Supplementary Figure 2A-D). We confirmed on sorted cells that Stx5 knock-down (KD) GFP^+^ PCs expressed less Stx5 at the transcriptional level than GFP^-^PCs (Supplementary Figure 2B). From two days after LPS-induced differentiation, PCs were generated from B cells and their frequency was roughly similar between control and Stx5 KD samples (Supplementary Figure 2C-D left panel). However, from day 4 onwards, the frequency of total PCs in the Stx5 KD samples progressively diminished to half of the frequency observed at day 2 while it was constant in the control samples during the same period (Supplementary Figure 2D left panel). Strikingly, the frequency of GFP^+^ PCs was rapidly decreasing in the Stx5 KD samples while it remains constant throughout the experiment in control samples, suggesting that Stx5 KD confers an intrinsic disadvantage to PCs (Supplementary Figure 2D, central panel). The frequency of GFP^+^ B cells was not modified by Stx5 KD (Supplementary Figure 2D, right panel), confirming the specific requirement of Stx5 expression for PC persistence *in vitro.* We next took advantage of a small molecule, Retro-2, reported to block Stx5 function by mislocalizing it and blocking its recycling^19,20^ to assess the impact of Stx5 on antibody secretion. When *in vitro* differentiated WT PCs were cultured for 5 hours in presence of increasing doses of Retro-2 we observed a dose dependent reduction of antibody secretion (Supplementary Figure 2E). Injection of a single dose of Retro-2 intraperitoneally to WT mice led to a significant reduction of the antibodies secreted by BM PCs after 3 days without reduction of the number of PCs in this organ (Supplementary Figure 2F), supporting a role for Stx5 in antibody secretion on top of PC survival. Altogether our data suggest that the machinery required for ER to Golgi transport including Stx5/Sec22b is upregulated during PC differentiation and that disruption of Stx5 leads to defects in PC viability and function.

### Dramatic loss of plasma cells and circulating antibodies in absence of Sec22b

Taking in consideration the effect of Stx5 on PC maintenance and antibody secretion, we next explored the implication of its partner Sec22b. We crossed a Sec22b floxed mouse model^11^ with the mb1-cre strain to generate a mouse model lacking Sec22b expression specifically in the B cell lineage (Sec22b^flox/flox^ x mb1-cre, hereafter referred to as Sec22b^B-KO^) (Supplementary Figure 3A-B). In absence of Sec22b, B cell development was roughly normal with only a mild reduction of the number of BM mature B cells and splenic follicular B cells compared to WT mice, whereas BM B cell precursors, splenic immature, marginal zone and CD93^-^/CD21^-^CD23^-^ B cells were unaffected (Supplementary Figure 3C-D). At steady state Sec22b^B-KO^ mice had almost no circulating antibody for all the isotypes tested (Figure 2A). Igκ representing around 90% of secreted antibodies in the mouse were reduced over 40 times. IgG1 titres were 55 times lower in absence of Sec22b. IgM, IgA and IgG3 were 40, 30 and 20 times lower in absence of Sec22b, respectively (Figure 2A).

**Figure 2:**
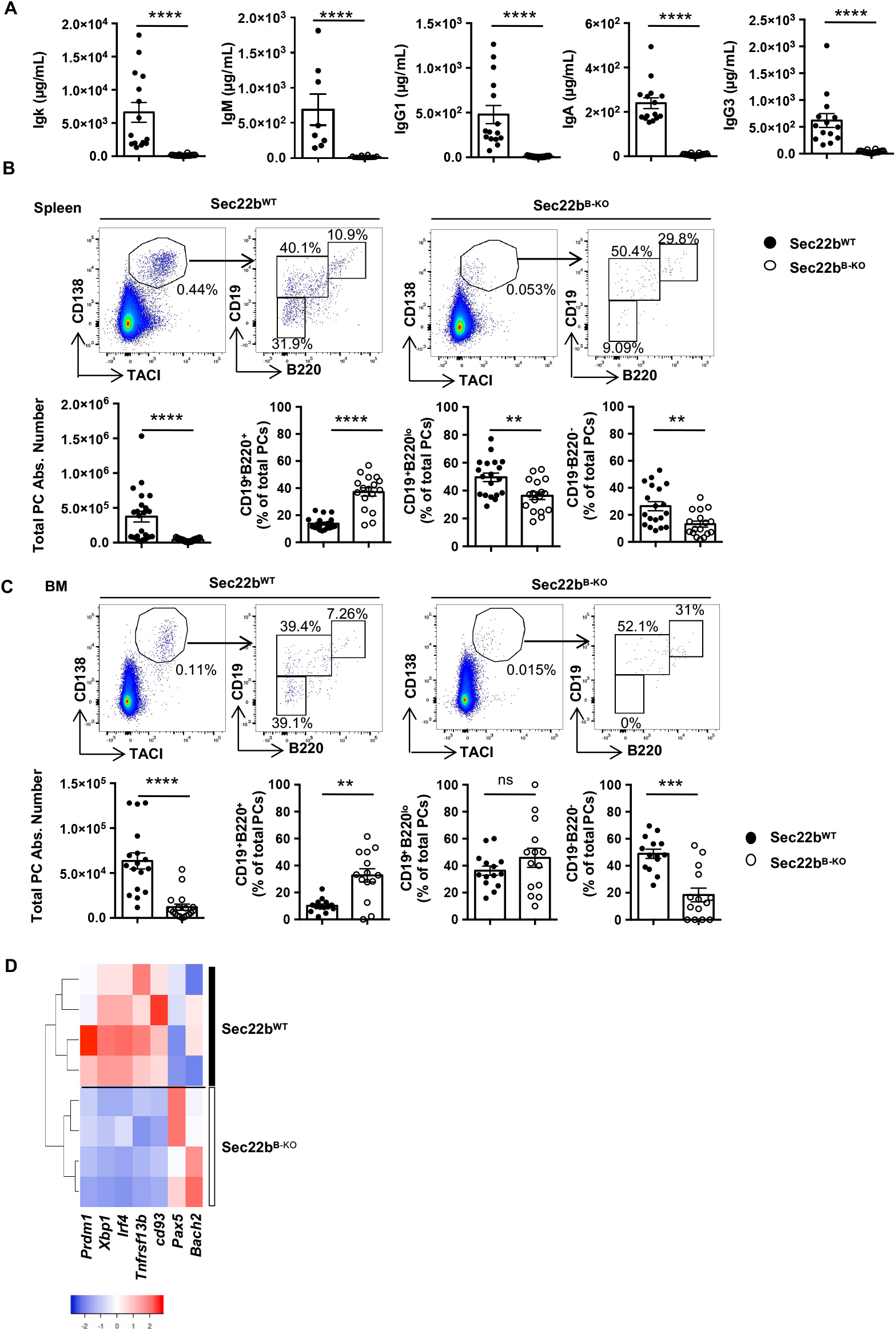
Dramatic loss of plasma cells and circulating antibodies in absence of Sec22b: A) ELISA quantification of Ig titres in sera of unimmunized Sec22b^WT^ or Sec22b^B-KO^ mice. B-C) Representative dot plots (top) and quantification (bottom) of absolute number of total PCs (CD138^+^TACI^+^) and percentage of each PC subsets: PBs (CD138^+^TACI^+^B220^+^CD19^+^), early PCs (CD138^+^TACI^+^B220^low^CD19^+^) and late PCs (CD138^+^TACI^+^B220^−^CD19^−^) in Sec22b^WT^ and Sec22b^B-KO^ mice in spleen (B) and bone marrow (C). Cells were first gated on their size, structure and viability. Dead cells and doublets were excluded. D) Unsupervised clustering based on the relative expression of *Prdm1, Xbp1, Irf4, Tnfrsf13b, CD93, Pax5 and bach2* of sorted Sec22b^WT^ and Sec22b^B-KO^ PCs determined by Biomark multiplex qPCRs at steady state. The heatmap was generated using the heatmapper.ca website and row Z score based on (2-^**D**Ct^) values. N= 7-19 mice from 2-5 independent experiments. The p-values were determined with the two-tailed Mann-Whitney non-parametric test. \**p* < 0.05; \*\**p* < 0.01; ***p <0.001 ****<p0.0001. “ns”= non-significant *p*-value.

This was associated with a ten-fold reduction of the frequency and absolute number of PCs in the spleen and the BM with the mature CD19^-^B220^-^ PCs being the most affected (Figure 2B and C). Accordingly, we observed that the remaining splenic PCs in Sec22b^B-KO^ mice display a less mature profile than their WT counterparts with reduced expression of the PC markers *Cd93* and *Tnfrsf13b* and of the PC master regulators *Prdm1, Xbp1* and *Irf4*. In contrast enhanced expression of the B cell master regulators *Pax5* and *Bach2* were detected at the transcriptional level in Sec22b^B-KO^ PCs compared to WT (Figure 2D). Thus, our results demonstrate that Sec22b plays a critical and non-redundant role in PC maintenance and in the control of antibody circulating titres in vivo.

### Sec22b is indispensable for the generation of a protective humoral immune response

Considering the importance of PCs and circulating antibodies for the humoral immune response, we next assessed the impact of *Sec22b* deficiency on these processes. Following T-dependent immunization with sheep red blood cells (SRBC) (Figure 3A-C), the frequency and number of PCs in the spleen remained extremely low in Sec22b^B-KO^ mice, being reduced over 100 times compared to controls (Figure 3B). A similar observation was made for antibody titres after SRBC immunization (Figure 3C). We also investigated antigen specific humoral immune response by immunizing with the T-dependent antigen NP-KLH in alum and boosting with NP-KLH only (Figure 3D). Seven days after the boost the frequency and numbers of splenic and BM PCs were again significantly reduced in absence of Sec22b (Figure 3E). Moreover, we barely detected NP-specific IgM and IgG1 antibodies in the serum of Sec22b^B-KO^ mice throughout the immunization while a potent antibody immune response with the expected kinetics was observed in control animals (Figure 3F). We next infected WT and *Sec22b* deficient mice with influenza A virus (Figure 3G) and observed a profound defect in the frequency and number of PCs in the draining mediastinal lymph nodes (Figure 3H). Flu-specific antibodies were undetectable in the serum of *Sec22b* deficient mice (Figure 3I). These defects were associated with exacerbated weight loss in Sec22b^B-KO^ mice compared to their WT littermates suggesting a poorer control of the infection in absence of Sec22b (Figure 3J). Altogether, these results establish that Sec22b is required for the establishment of a potent and efficient humoral immune response after both vaccination and infection.

**Figure 3:**
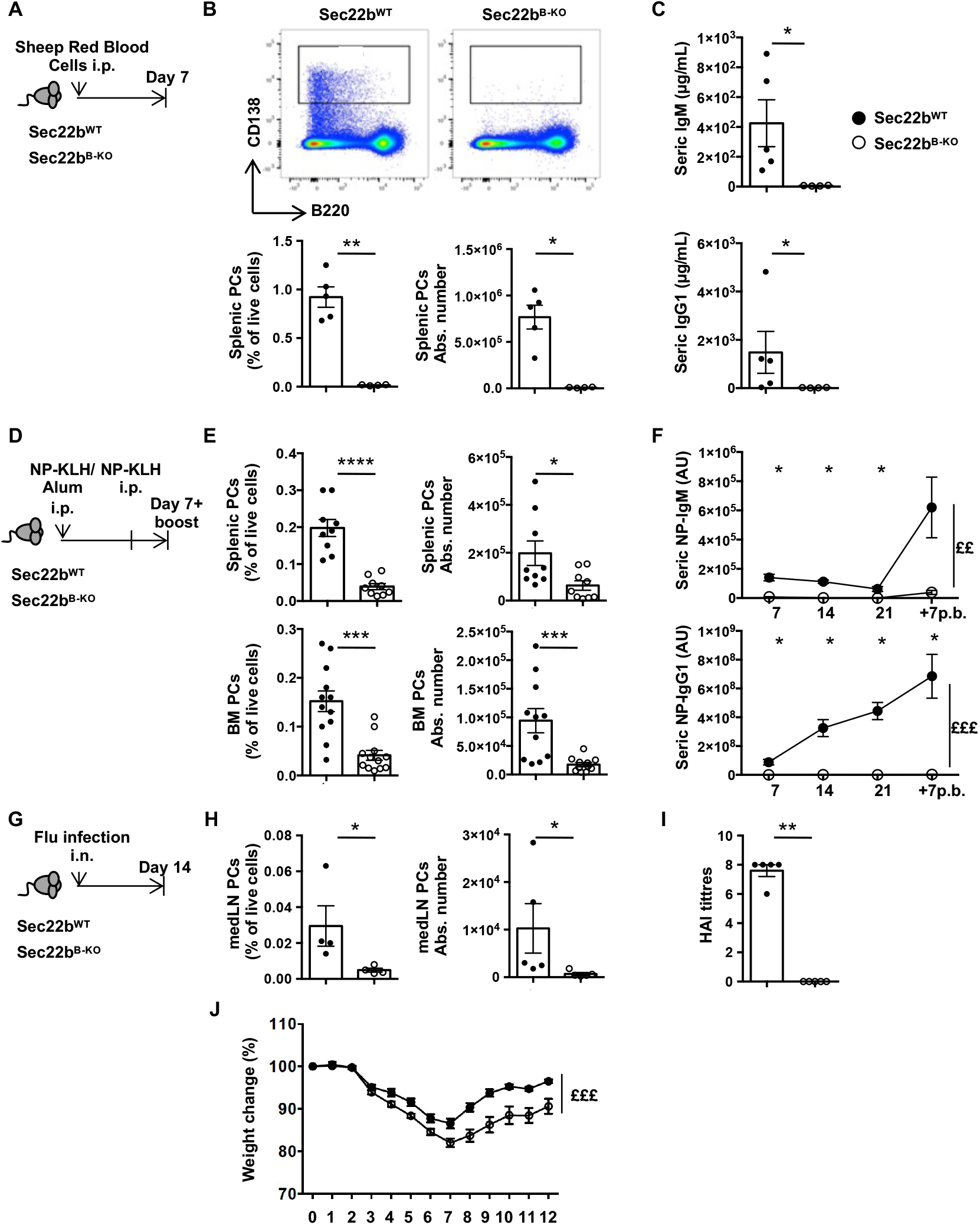
Sec22b is indispensable for the generation of a protective humoral immune response: A) Schematic representation of the SRBC immunisation protocol for Sec22b^WT^ and Sec22b^B-KO^ mice. B) Representative dot plots (top), frequency (bottom left) and absolute number (bottom right) of splenic PCs (CD138^+^B220^+/-^ determined by flow cytometry 7 days after SRBC immunization. C) ELISA quantification of IgM (top) and IgG1 (bottom) serum titres 7 days after SRBC immunized. A-C one representative experiment of 2 is shown. D) Schematic representation of the NP-KLH immunization/ boost protocol. Sec22b^WT^ or Sec22b^B-KO^ mice were immunized, rechallenged 28 days later and analysed 7 days later. Sera were collected at days 7-, 14-, 21- and 7-days post boost. E) Frequency (left) and absolute number (right) of splenic PCs (CD138^+^B220^+/-^) (top) and bone marrow (BM) PCs (CD138^+^B220^+/-^) (bottom) determined by flow cytometry 7 days post boost with NP-KLH (n=9 mice from 2 pooled independent experiments). F) ELISA quantification of NP-IgM (top) and NP-IgG1 (bottom) serum titres at days 7-, 14-, 21-post primary immunization and 7-days post boost (n=5 mice. One experiment representative of 2 is shown). G) Schematic representation of the Flu infection protocol for Sec22b^WT^ and Sec22b^B-KO^ mice. Mice were analysed 14 days after infection with influenza A virus. H) Frequency (left) and absolute number (right) of mediastinal PCs (CD138^+^B220^+/-^) determined by flow cytometry 14 days after infection with influenza A virus. I) Quantification of HAI titres 14 days post flu infection. J) Weight change of Sec22b^WT^ or Sec22b^B-KO^ mice over 12 days after flu infection. (H-I; n=5, one representative experiment of 2 is shown, J; n=15 from 2 pooled experiments) For flow cytometry experiment cells were first gated on their size and structure, then their viability. Dead cells and doublets were excluded. The p-values were determined with the two-tailed Mann-Whitney non-parametric test \**p* < 0.05; \*\**p* < 0.01; ***p <0.001; ****<p0.0001 or with the 2way ANOVA with Sidak correction for multiple comparisons (**££**p<0.01, **£££**p<0.001).

### Sec22b is necessary for plasma cell maintenance and secretory function but not for the initiation of differentiation

To unravel at which step of PC differentiation Sec22b is required, we performed *in vitro* differentiation assays of splenic B cells. In control cultures PC frequency doubles between day 2 and day 4, whereas it remains constant between these two time points in Sec22b^B-KO^ cultures (Figure 4A-B).

**Figure 4:**
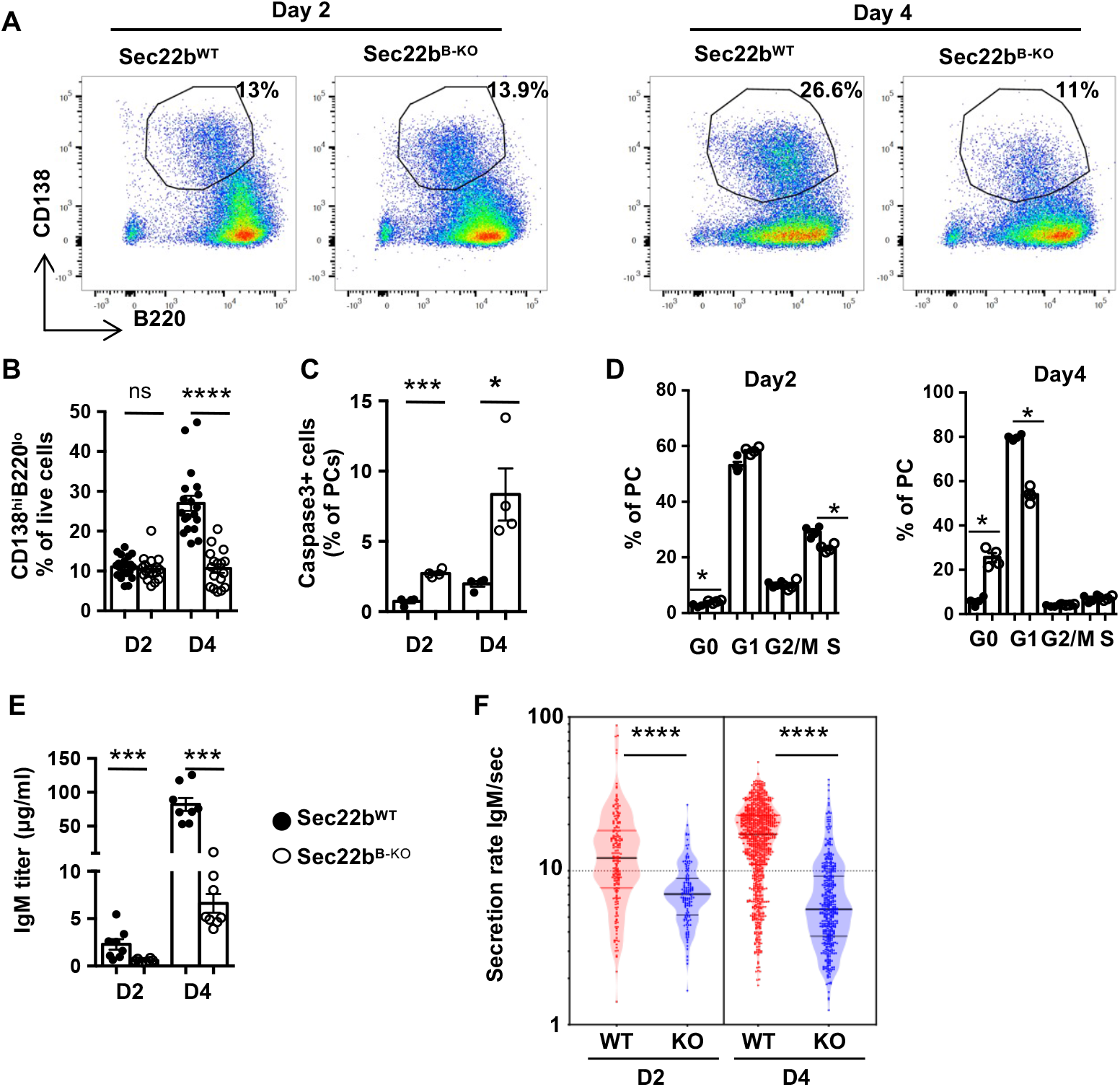
Sec22b is necessary for plasma cell maintenance and function. A-B) Representative dot plots (A) and quantification (B) of PCs (CD138^+^B220^+/-^) generated *in vitro* from Sec22b^WT^ and Sec22b^B-KO^ splenocytes after 2 or 4 days of stimulation with LPS. Cells were gated on their size and structure, on their viability and doublets were excluded. C) Frequency of Caspase 3^+^ PCs at day 2 and 4 post LPS stimulation determined by flow cytometry. D) Flow cytometry analysis of the cell cycle phases based on DAPI and Ki-67 staining in *in vitro* generated Sec22b^WT^ and Sec22b^B-KO^ PCs at day 2 (left) and day 4 (right) post LPS stimulation. E) ELISA quantification of total IgM secreted in the culture supernatant of PCs generated from Sec22b^WT^ and Sec22b^B-KO^ splenocytes after 2 or 4 days of LPS stimulation. F) Quantification of the IgM secretion rate from *in vitro* generated PCs from Sec22b^WT^ and Sec22b^B-KO^ splenocytes after 2 or 4 days of LPS stimulation analysed by DropMap. Each point represent one cell. (A-B) n=12-16 mice from at least 3 pooled independent experiments. (C-D) n=4 of 1 representative experiment out of 2. (E) n=8 in 3 independent experiments; (F) n=2 in 2 independent experiments. The p-values were determined with the two-tailed Mann-Whitney non-parametric test \**p* < 0.05; \*\**p* < 0.01; ***p <0.001, “ns”= non-significant *p*-value.

This was associated with an enhanced apoptosis of *in vitro* derived PCs lacking Sec22b as detected by the frequency of active Caspase3^+^ cells (Figure 4C). In addition, cell cycle was impaired in Sec22b^B-KO^ PCs generated *in vitro* with a progressive exit from the cell cycle particularly clear at day 4 (Figure 4D). These results indicate that Sec22b is not required for the initiation of PC differentiation but is important for the maintenance of this cell subset. In line with the reduced frequency of PCs generated we observed a decreased amount of total IgM in the culture supernatant in absence of Sec22b (Figure 4E). To assess more precisely the impact of *Sec22b* deficiency on antibody secretion we took advantage of a droplet microfluidic-based technique to assess the secretion rate at the single cell level ^1^. Between 6000 and 10000 cells were individually encapsulated into droplets and analysed for their IgM secretion after 2- and 4-days of *in vitro* culture. Among them up to 12% secreted detectable amount of IgM over the 40 minutes of imaging with less secreting cells in the Sec22b^B-KO^ condition as expected. We observed that *in vitro* generated WT PCs secreted on average 12 IgM/sec at day 2 and 20 IgM/sec at day 4. In contrast, *Sec22b* deficient PCs secreted on average 2 to 4 times less IgM/sec at day 2 and 4, respectively (Figure 4F). Thus, *Sec22b* deficiency leads to a significant reduction, albeit not a total block, in antibody secretion and to a dramatic reduction of PC survival and proliferation.

### Sec22b deficiency profoundly alters the PC transcriptome

To gain further insight into the molecular mechanisms under the control of Sec22b we performed RNAseq analysis on *in vitro* generated PCs. We chose an early time point, 2 days after LPS stimulation, to unravel defective mechanisms before the loss of PCs observed rapidly thereafter. Despite being numerically normal (Figure 4A-B), *Sec22b* deficient PCs at day 2 post stimulation were transcriptionally very distinct from their WT counterparts as shown by unsupervised analyses (Figure 5A-B and Supplementary Tables 1 and 2). Over 6000 genes were differentially regulated between both genotypes with 3388 genes up-regulated and 3184 genes down-regulated in Sec22b^B-KO^ PCs compared to WT PCs (14026 genes total, fold change >1.2 and q-value < 0.05) (Figure 5C). Gene set enrichment analyses (GSEA) revealed that several pathways were significantly different between WT and Sec22b^B-KO^ PCs. Pathways pertaining to “Cell cycle/mitosis” but also to “Mitochondria” and “Myc targets” were significantly downregulated in Sec22b^B-KO^ compared to WT PCs, whereas “UPR”, “ER-Golgi transport” and “protein secretion” pathways were upregulated (Figure 5D and Supplementary Figures 4 and 5).

**Figure 5:**
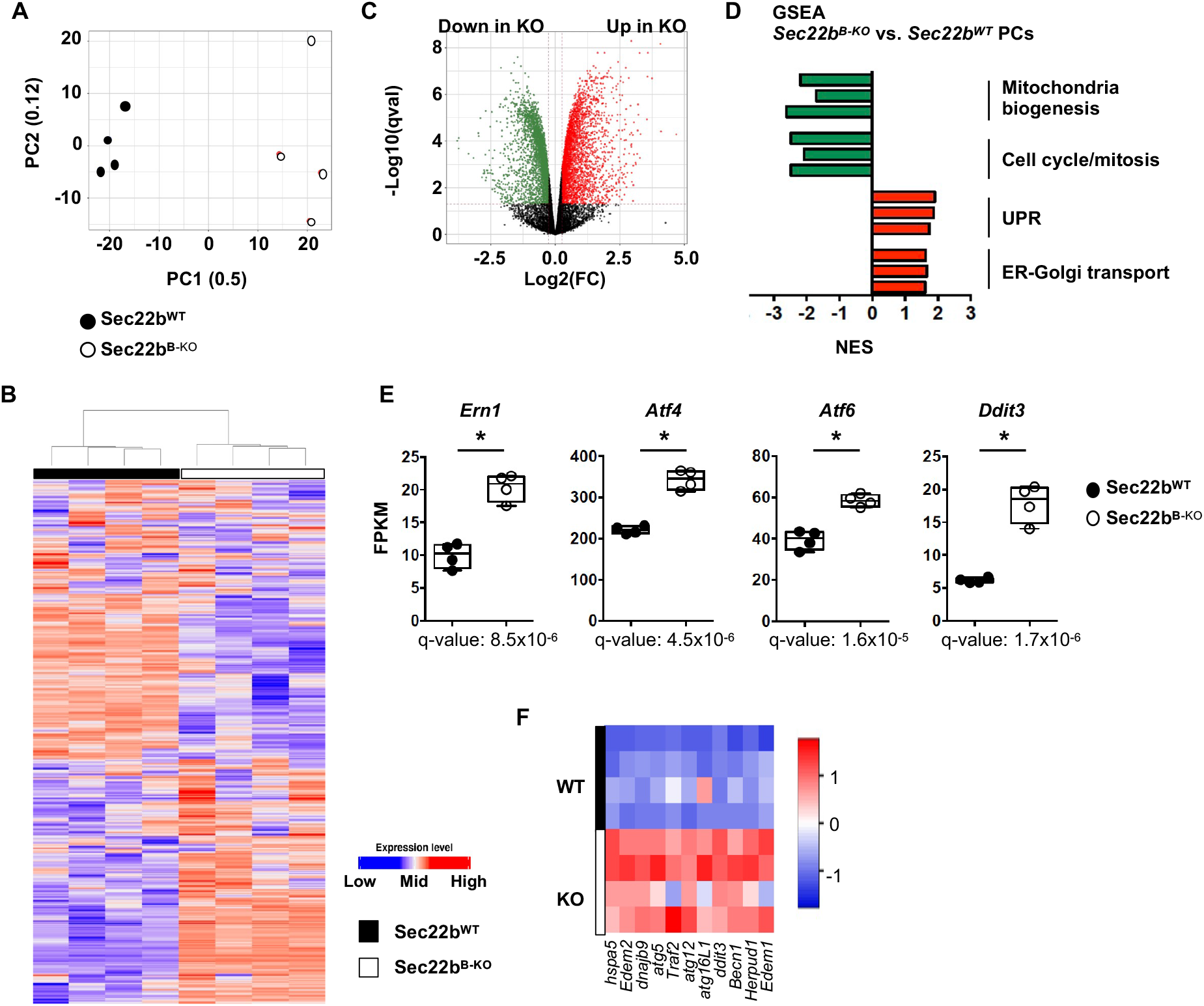
Sec22b deficiency profoundly alters the PC transcriptome. (A-E) RNAseq analysis of Sec22b^WT^ and Sec22b^B-KO^ PCs generated *in vitro* at day2 post LPS stimulation. A) Principal component analysis of the RNAseq data of Sec22b^WT^ and Sec22b^B-KO^ PCs. The proportion of variance is indicated for PC1 and PC2. B) Unsupervised clustering of Sec22b^WT^ and Sec22b^B-KO^ PCs based on the 500 most differentially expressed genes. C) Volcano-plot showing the differentially expressed genes between Sec22b^WT^ and Sec22b^B-KO^ PCs. Genes significantly downregulated and upregulated in Sec22b^B-KO^ PCs are shown in green and red, respectively (FC>1.2; q value<0.05). D) Normalized Enrichment Scores (NES) of representative gene sets significantly enriched in Sec22b^B-KO^ vs Sec22b^WT^ PCs and characteristic of selected cellular pathways. Gene sets significantly downregulated and upregulated in Sec22b^B-KO^ PCs are shown in green and red, respectively. E) Expression of *Ern1* (encoding Ire1a), *Atf4, Atf6, Ddit3* (encoding Chop) in fragments per kilobase of exon per million reads mapped (FPKM) determined by RNAseq in Sec22b^WT^ and Sec22b^B-KO^ PCs. F) Heatmap showing the relative expression of selected ER stress genes from PCs generated from Sec22b^WT^ and Sec22b^B-KO^ splenocytes after 2 days of LPS stimulation, determined by Biomark multiplex qPCRs at steady state. The heatmap were generated using the heatmapper.ca website, row Z score based on (2-^**D**Ct^) values. N=4 and data are representative of 1 or 2 (F) experiments. The p-values were determined with the two-tailed Mann-Whitney non-parametric test \**p* < 0.05.

These results suggest that *Sec22b* expression is key for the regulation of PC biology and highlight several Sec22b-dependent pathways. In line with our experimental results (Figure 4D), Sec22b seems essential for promoting cell cycle possibly via control of the Myc signalling axis. In absence of Sec22b, genes involved in ER-Golgi transport and secretion were increased despite the observed reduced antibody secretion rate. This could correspond to a compensatory mechanism to cope with defective constitutive secretion. Accordingly, and representing a clear sign of exacerbated ER stress, all three branches of the UPR (i.e. *Ern1, Atf4/Ddit3, Perk, Atf6*) (Figure 5E-F) were upregulated in Sec22b^B-KO^ PCs including the Atf4/Chop pathway normally actively repressed in PCs to protect from ER-stress induced cell death^4^. Our results suggest that in absence of Sec22b this protective mechanism is deficient leading to Chop-mediated cell death and exit from the cell cycle. Furthermore, our RNAseq data suggests that *Seb22b* deficiency may contribute to defects in several organelles including the ER and Golgi apparatus.

### Sec22b is essential for ER expansion and structure in PCs

As genes involved in ER-Golgi transport were up-regulated in Sec22b^B-KO^ (Figure 5D-F) mice while the secretion was reduced (Figure 4E-F) we explored more precisely how *Sec22b* deficiency may affect these two organelles. We observed using a permeable cell tracker that the ER expansion normally observed as PC differentiate was significantly reduced in absence of Sec22b (Figure 6A). The structure of the ER was also altered in Sec22b^B-KO^ PCs. Whereas the perinuclear envelope was roughly normal, peripheral ER appeared poorly branched with dilated cisternae filled with IgM (Figure 6B), suggestive of accumulation of hyper-dilated ER and loss of the parallel rough ER typically seen in WT PCs. Electron microscopy confirmed that the stacked ER sheets characteristic of PCs were lost in absence of Sec22b with a hyper-dilatation of the ER cisternae and a defective stacking that was even more evident at day 4 (Figure 6C).

**Figure 6:**
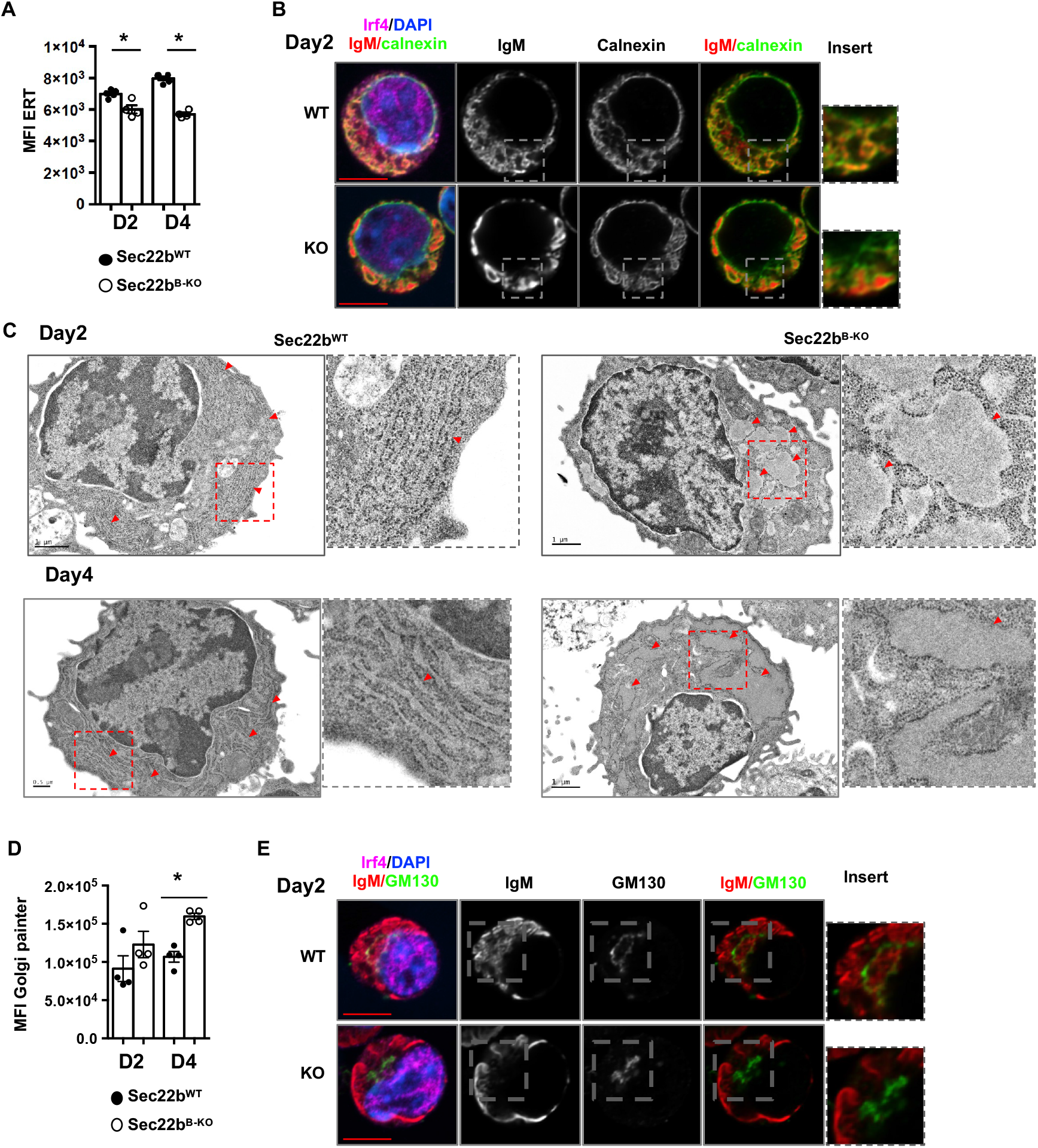
Sec22b is essential for ER expansion and structure in PCs. A) Flow cytometric quantification of ER-Tracker MFI (geometrical mean) on PCs generated from Sec22b^WT^ and Sec22b^B-KO^ splenocytes after 2 or 4 days of LPS stimulation. B) Confocal microscopy images of Sec22b^WT^ and Sec22b^B-KO^ PCs obtained from splenocytes stimulated with LPS for 2 days. Cells were stained with anti-IgM to detect antibodies, anti-calnexin to detect the ER and anti-IRF4 antibodies. Nuclei were counterstained with Hoescht. C) Electron microscopy images of Sec22b^WT^ (left) and Sec22b^B-KO^ (right) PCs obtained from splenocytes stimulated with LPS for 2 (top) or 4 (bottom) days. Red arrowheads indicate ER sheets. Scale bar = 1mm or 0.5mm. D) Flow cytometric quantification of Golgi painter MFI (geometrical mean) on PCs generated from Sec22b^WT^ and Sec22b^B-KO^ splenocytes after 2 or 4 days of LPS. E) Confocal microscopy images of Sec22b^WT^ and Sec22b^B-KO^ PCs obtained from splenocytes stimulated with LPS for 2 days. Cells were stained with anti-IgM, anti-GM130 to detect the Golgi and anti-IRF4 antibodies. Nuclei were counterstained with Hoescht. N=4 of 1 representative experiment out of 3. The p-values were determined with the two-tailed Mann-Whitney non-parametric test \**p* < 0.05.

This suggests that Sec22b expression is crucial for ER spatial expansion and organisation during PC differentiation. In the conventional constitutive secretion route that antibodies were reported to use, proteins leave the ER through the Endoplasmic Reticulum-Golgi Intermediate Compartment (ERGIC) to reach the Golgi apparatus where their final glycosylation occurs. The Golgi apparatus appeared perturbed in Sec22^B-KO^ PCs with enhanced intensity of the permeable tracker Golgi painter at day 4 (Figure 6D) and reduced detection of colocalized IgM and of the Golgi marker GM130 (Figure 6E), suggestive of a block of antibody trafficking between the ER and the Golgi. Altogether, our results imply a critical role for Sec22b as a regulator of ER organization, structure and function in PCs with repercussion on the Golgi apparatus as well.

### Sec22b deficiency affects PC fitness via altered mitochondrial dynamics

In addition to protein folding and export, an important function of the ER is the control of organelle dynamics and in particular of mitochondria. Indeed, ER-mitochondria membrane contact sites are pivotal for mitochondrial fission and thus function^21^. Through our RNAseq analysis we observed a downregulation of several genes involved in mitochondrial function and dynamics (Figure 5D and Supplementary Figure 4) including *Drp1, Inf2* and *Spire1* that encode proteins involved in mitochondrial fission at the ER-mitochondria contact site (Figure 7A). Mitochondrial content and potential were indeed significantly increased in PCs after 4 days in culture in Sec22b^B-KO^ PCs compared to WT PCs (Figure 7B). Accordingly, we detected more mitochondria, increased total mitochondrial area but with fewer fragments in Sec22b^B-KO^ PCs compared to WT PCs (Figure 7C) suggesting an hyperfused mitochondrial phenotype in Sec22b^B-KO^ PCs (Figure 7D). We wondered whether this defective mitochondrial conformation may contribute to the PC loss observed both *in vivo* and *in vitro* in absence of Sec22b. Treatment of WT PCs with M1, a Drp1 antagonist, together with Mdivi, an agonist of mitochondrial fusion, promoted the generation of a hyperfused mitochondria phenotype comparable to that of Sec22b^B-KO^ PCs (Figure 7E). Moreover, M1+Mdivi-treated WT PCs numbers were significantly reduced in culture compared to control conditions, confirming that mitochondrial dynamics affect PC fitness (Figure 7F). In addition to this hyperfused phenotype Sec22b^B-KO^ PCs presented several abnormalities in mitochondrial gene expression (Figure 7G) including genes implicated in the respiratory chain (*Atp5* family), mitochondrial fission (*Mtfp1* and *Mtfr2*), structure (*Timm* and *Tomm* families) and metabolite transport (*Slc25* family) (Figure 7G) suggesting a global alteration of mitochondrial size and function in absence of Sec22b. Altogether, these results suggest that Sec22b expression regulates mitochondrial dynamics in PCs with important consequences for cell fitness.

**Figure 7:**
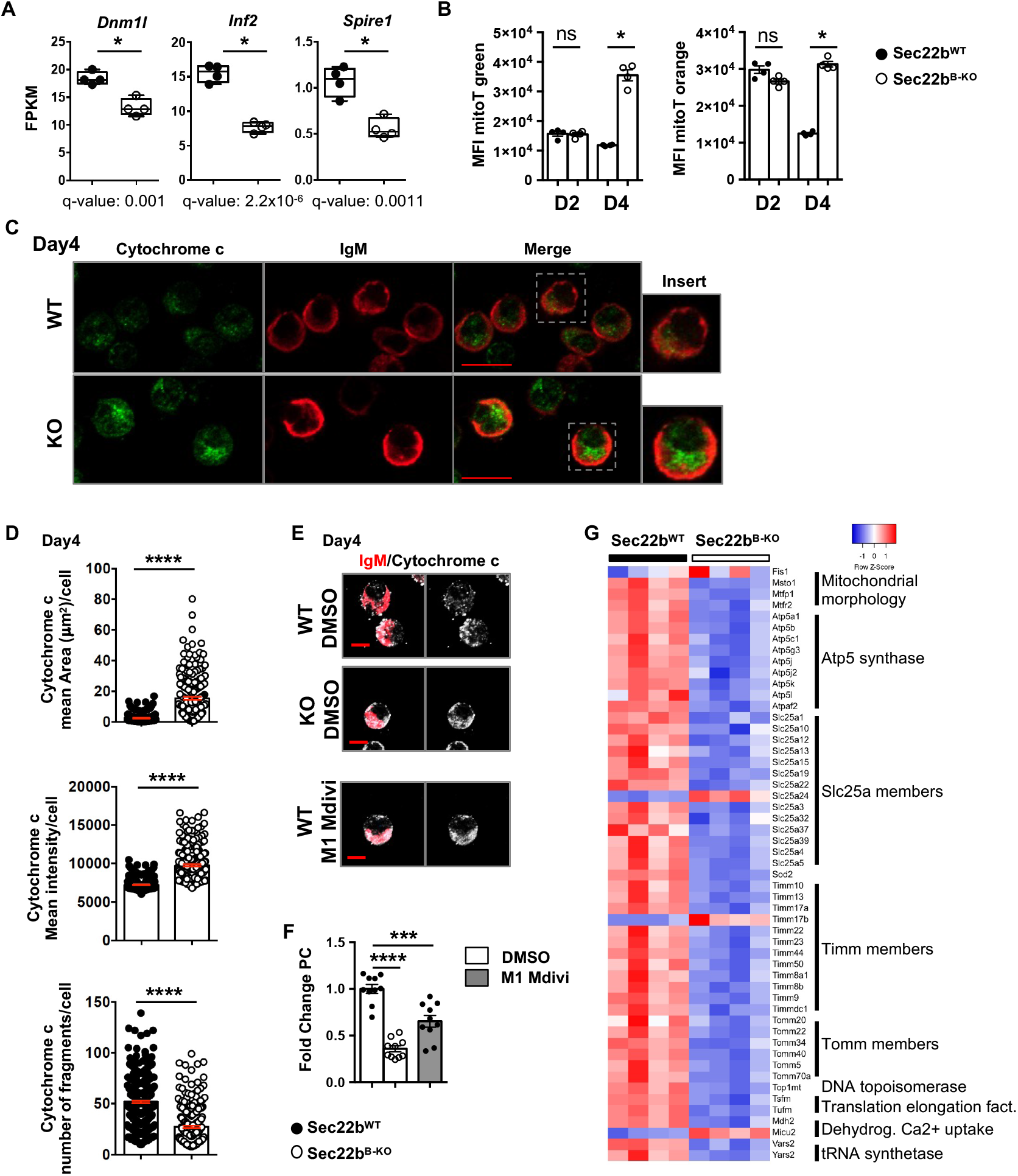
Sec22b deficiency affects PC fitness via altered mitochondrial dynamics. A) Expression of *Dnm1l, Inf2* and *Spire1* in FPKM determined by RNAseq in Sec22b^WT^ and Sec22b^B-KO^ PCs obtained from splenocytes stimulated with LPS for 2 days. n=4 and data are representative of 1 experiment B) Flow cytometric quantification of Mito-tracker (mitoT) green (left) and mitoT orange (right) MFI (geometrical mean) on PCs generated from Sec22^WT^ and Sec22b^B-KO^ splenocytes after 2 or 4 days of LPS stimulation. n= 4 mice in 1 representative experiment out of 3. C-D) Confocal microscopy images of PCs obtained from Sec22b^WT^ and Sec22b^B-KO^ splenocytes stimulated with LPS for 4 days. Cells were stained with anti-IgM to detect antibodies, anti-cytochrome c to detect mitochondria, anti-IRF4 antibodies and nuclei were counterstained with Hoescht. Scale bar = 10mm. Representative images (C) and quantification (D) of the mean area (top), mean intensity (mean per z-stack) (middle) and number of fragments (over the full cell volume quantified through 10 z-stacks) (bottom) per cell of the cytochrome c staining in PCs obtained from Sec22^WT^ and Sec22b^B-KO^ splenocytes stimulated with LPS for 4 days. n=304 for the WT and n=231 for the Sec22b^B-KO^ from 3 independent mice per genotype. E-F) 3D projection of confocal microscopy images of PCs obtained from Sec22^WT^ and Sec22b^B-KO^ splenocytes stimulated with LPS for 4 days and after 2 days of treatment with DMSO or M1/Mdivi for Sec22^WT^. Cells were stained with anti-IgM, anti-cytochrome c antibodies and nuclei were counterstained with Hoescht. Scale bar = 6mm. Representative images (E) and quantification (F) of fold change of PC frequency normalized to control Sec22b^WT^ PCs are shown. n=10 from 2 pooled experiments. G) Supervised heatmap based on RNaseq expression data of 9 mitochondrial gene families differentially expressed between Sec22b^WT^ and Sec22b^B-KO^ PCs. For flow cytometry experiment cells were gated on their size and structure, on their viability and doublets were excluded. The p-values were determined with the two-tailed unpaired Mann-Whitney non-parametric test \**p* < 0.05; ****p <0.0001, “ns” = non-significant *p*-value.

## Discussion

Despite their essential role in health and disease, how PCs maintain a high rate of antibody secretion and survive this massive protein synthesis and processing burden is unclear. Here we demonstrate that SNAREs, and in particular Sec22b, are essential for these processes. Indeed, in absence of *Sec22b* expression in the B cell lineage, the humoral immune response is severely impaired due to decrease in antibody secretion and defective PC maintenance.

The reduced antibody secretion rate observed is likely caused by a defective ER to Golgi transport due to altered formation of the Stx5/Sec22b SNARE complex. Indeed, fewer IgM were detected in the Golgi apparatus in absence of Sec22b while large aggregates were observed in the dilated ER. Supporting a role for this complex in this process, PC treatment with Retro-2, a drug known to block Stx5 canonical function^19^, also leads to decreased antibody secretion. Interestingly, after Retro-2 treatment but also in absence of Sec22b, antibody secretion was reduced but not totally abrogated suggesting that redundant mechanisms may exist to export antibodies via the classical constitutive or via unconventional secretion pathways^22,23^.

The altered PC maintenance in absence of Sec22b cannot uniquely be linked to the defective secretory capacity of the cells as several studies report that PC survival and antibody secretion can be uncoupled. For example, mice lacking Xbp1 or the ligase Rctb, important for Xbp1 activation, display reduced antibody secretion but normal PC numbers^24–27^. Moreover, PC producing pathogenic antibodies often fail to secrete them while the PC compartment is normal in these mice^28^. Moreover PCs generated in the LMP2A model can survive without secreting any antibody demonstrating the independence between antibody secretion and PC survival^29^. Additive defects must thus explain the poor persistence of PCs in absence of sec22b. Importantly, we report here that *Sec22b* deficiency is associated with deregulation of the UPR as well as morphological alterations of the ER and mitochondria.

In PCs, the Ire1α/Xbp1 axis is normally highly expressed and favours antibody production and folding without causing cell death^2,30^ whereas Atf6 is weakly expressed and the Perk-dependent branch of the UPR is repressed^4,31^. Activation of the Perk/Atf4/Chop pathway has been associated with increased susceptibility to cell death in PCs and is normally suppressed by the Ire1α/Xbp1-dependent Ufbp1 protein^32^. In absence of Sec22b the up-regulation of the three UPR branches may have several consequences. The upregulation of Ire1α may contribute to compensatory mechanisms to deal with the decreased ER to Golgi protein transport and for sustaining sub-optimal antibody secretion. In parallel, the upregulation of Perk may lead to the inhibition of the cell cycle we observed^33^ and eventually to PC cell death^34^ hence explaining, in part, the poor PC maintenance *in vitro* and *in vivo* in absence of Sec22b. Another important observation made in absence of Sec22b was the disturbed ER structure with poorly stacked and hyper-dilated cisternae that contain antibody aggregates. This could be the consequence of the reduced transport of antibodies from the ER to the Golgi apparatus but could also be caused directly by the absence of Sec22b as was shown in yeast and drosophila^35–37^. In yeast, Sec22p interacts with Sey1p to regulate sterol biosynthesis and with atlastins to support ER homotypic fusion^35,36^. In the drosophila, Sec22 was identified as an important regulator of ER structure with a profound expansion of the ER when Sec22 was mutated or knocked-down^37^. More recently, Sec22b knock-down in HUVECs was also associated with dilated ER but the mechanism at play remain unclear^38^. Interestingly, the long form of Stx5 was also shown to be important for ER branching via interaction with Climp63 that links the ER membrane to the microtubules^14^. Thus, Sec22b could directly control ER morphology in PCs either by interacting with the long form of Stx5 or via another partner still to identify.

Another striking defect observed in *Sec22b* deficient PCs was the accumulation of hyperfused mitochondria. As previously shown in other cell types this could be a consequence of the pronounced cell cycle exit, as observed herein in Sec22b^B-KO^ PCs. Indeed, the kinase CDK1 that is strongly downregulated in absence of Sec22b phosphorylates Drp1 to promote its recruitment to mitochondria and the induction of mitochondrial fission^39^. Alternatively, the disrupted structure of the ER in absence of Sec22b may alter ER/mitochondria contact sites that regulate mitochondrial fission^21^. Sec22b, through its longin domain, has already been shown to be important for ER/plasma membrane contact sites and consequently for plasma membrane expansion in neurites^15,40^. Whether Sec22b also contributes to ER/mitochondria contact sites directly or indirectly remains to be established. In addition, we demonstrated that hyperfused mitochondria were associated with poor PC survival hence suggesting that altered mitochondrial dynamics contribute to the decreased PC number in absence of Sec22b.

To conclude, we demonstrated that Sec22b is a crucial regulator of antibody secretion by regulating transport between the ER and the Golgi apparatus, a key bottleneck in this process. Moreover, through regulation of the UPR pathways and ER/mitochondrial dynamics, Sec22b is paramount for PC survival. Altogether, our results thus demonstrate that Sec22b-mediated regulation of organelle dynamics is indispensable for PC biology and for the establishment of a protective humoral immune response. They also open new avenues for targeting PC in pathological contexts.

## Acknowledgments

We thank Dr. N. Setterblad, C. Doliger and S. Duchez (Plateforme technologique IRSL, Paris, France) and Dr. V. Parietti (Mouse facility IRSL, Paris, France) for their technical assistance. The study was supported by the Laboratory of Excellence in Research on Medication and Innovative Therapeutics (LabEx LERMIT) (ME and KB), an ANR JCJC grant (ANR-19-CE15-0019-01), an ANR @RAction grant (ANR-14-ACHN-0008), a “Fondation ARC pour la recherche sur le cancer” grant (P JA20181208173) and a grant from IdEx Université de Paris (ANR-18-IDEX-0001) to ME, an ANR PRC grant (ANR-17-CE14-0019) and an INCa grant (PRT-K 2017) to KB. P.B. acknowledges funding from the French National Research Agency grant ANR-18-CE15-0001 project Autoimmuni-B, by the Institut Carnot Pasteur Microbes et Santé grant ANR-11-CARN-0017-01, the Institut Pasteur and the Institut National de la Santé et de la Recherche Médicale (INSERM). M.A.L is supported by Biotechnology and Biological Sciences Research Council (BBS/E/B/000C0427, BBS/E/B/000C0428, and the Campus Capability Core Grant to the Babraham Institute). AP and MC were supported from a grant from the Biotechnology and Biological Sciences Research Council (BB/L022389/1). DLH is supported by a National Health and Medical Research Council Australia Early-Career Fellowship (APP1139911). NA was supported by a PhD fellowship from the French Ministry for education and by a 4^th^ year PhD fellowship from the “Fondation ARC pour la recherche sur le cancer”. P.C-H. was supported partly by a stipend from the Pasteur - Paris University (PPU) International PhD program, and by a fellowship from the French *Fondation pour la Recherche Médicale* (FRM). The “EMiLy” U1160 INSERM unit is a member of the OPALE Carnot institute, The Organization for Partnerships in Leukemia (Institut Carnot OPALE, Institut de Recherche Saint-Louis, Hôpital Saint-Louis, Paris, France. Web : www.opale.org.).

## Author Contributions

AB designed and performed experiments, analyzed data, and wrote the manuscript. LG, SMG, ER, PCH, DS, DLH, NR, MC and MAL performed experiments and analyzed data. NA helped with experiments. SB and VS performed electron microscopy experiments. AA, DG, JB, PB and SA provided essential reagents and technologies. KGCS and FB contributed to the project design. KB, MAL and AAP contributed to the project design and to the manuscript redaction. ME designed the project, analyzed data, and wrote the manuscript. All authors had the opportunity to review and edit the manuscript.

## Declaration of Interests

The authors declare that no conflict of interest exists.

## Methods

### Mouse model, immunisation, and infection

The Sec22b^fl/fl^ mice (C57Bl/6J background) were obtained from Dr. Sebastian Amigorena and crossed with the mb1-Cre mice (C57Bl/6J background) purchased from The Jackson Laboratory. All experiments were conducted in compliance with the European Union guide for the care and use of laboratory animals and has been reviewed and approved by appropriate institutional review committees (C2EA-26, Animal Care and Use Committee, Villejuif, France and Comité d’éthique Paris-Nord N°121, Paris, France). Immunizations/infections were performed intraperitoneally (ip) with 100μg of 4-hydroxy-3-nitrophenylaceyl-keyhole limpet hemocyanine (NP-KLH) (Biosearch Technologies) adjuvanted with alum (Inject Alum, Thermo Scientific) or with 200μL of Sheep Red Blood Cells (SRBC) (Eurobio) or intranasally with 10^4^ plaque-forming units of influenza A/HK/x31 virus (H3N2) under inhalation anaesthesia with isoflurane. Influenza infections were performed on chimeric mice. CD45.1 Recipient mice were lethally irradiated with 11gy before reconstitution with bone marrow cells from Sec22b^B-KO^ or Sec22b^B-WT^ mice. Chimera were infected 8 to 12 weeks later once reconstitution was complete.

### Cell preparation and flow cytometry

Spleen and bone marrow cells were isolated as previously described^41,42^. Briefly, single cell suspensions were stained with appropriate antibodies (Supplementary table 3) in PBS supplemented with 2%BSA and 2mM EDTA for cell surface staining. Staining with ERtracker (Invitrogen), mitoTracker Green (Invitrogen) and mitoTracker Orange (Invitrogen) were performed as recommended by the supplier. Intracellular staining was performed using the FoxP3/Transcription factor staining buffer set (eBioscience) according to the provider recommendation. Flow cytometry analyses were performed on a BD LSR Fortessa cytometer and cell sorting experiments for RNAseq and Biomark analysis were performed using a BD FACS AriaIII cell sorter. Data were analysed with the Flowjo software (TreeStar, Ashland, OR).

### In vitro cell differentiation and infection of primary cells

1<10^6^ splenocytes were stimulated with 5μg/mL of lipopolysaccharide (LPS) (InVivoGen) for 2 or 4 days in complete culture medium composed of RPMI supplemented with 10% foetal calf serum (Sigma), 0.05mM 2-mercaptoethanol, 100U/mL penicillin-streptomycin, 1mM sodium pyruvate and 0.1mM nonessential amino acids (Gibco). Where indicated, cells were treated with 20μM M1 and 10μM Mdivi (Sigma) for 2 days from day 2 to day 4.

*Stx5a* specific shRNAs were designed thanks to the RNAi consortium (https://portals.broadinstitute.org/gpp/public) and cloned in the pLKO.3G (addgene #14748) vectors. Lentiviral particles were produced in the Human Embryonic Kidney 293T (HEK293T) cell line with the psPAX2 (Addgene #12260) and pMD2.G (Addgene #12259) vectors. For primary cell transduction, splenocytes were put in culture in complete culture medium in presence of 80ng/ml CD40L (Thermoscientific) and 1U/mL IL-4 (Miltenyi) for 24 hours to promote B cell entry into cycle. The cells were then washed and transduced with lentiviral particles together with polybrene. After 24hours cells were washed and differentiated into PCs by addition of LPS as indicated above.

### Western blot

Cells were resuspended in RIPA lysis buffer supplemented with Protease and Phosphatase Inhibitor (Thermofisher). Total proteins were quantified with Bradford buffer (Thermofisher). 15μg of proteins were separated on a NuPAGE™ 4-12% Bis-Tris Gel (Invitrogen) and transferred to a PVDF membrane. Primary antibodies (Supplementary Table 3) or β-actin (Cell Signaling) were incubated overnight at 4°C. Secondary antibodies (Supplementary Table 3), specific for the primary antibody species, conjugated to horseradish peroxidase (HRP) were incubated 2h at room temperature and detected using Pierce ECL (Thermofisher) and signal was quantified by ChemiDoc™ Touch Gel Imaging System (BIO RAD).

### Confocal microscopy

Cells were loaded on poly-Lysine (Sigma) coated slides and fixed in PBS/4% PFA prior to analysis. Cells were permeabilized with PBS/0.3% Triton, washed in PBS and incubated with primary antibodies coupled or not overnight at 4°C and then with secondary antibodies 1 hour at room temperature, when necessary (Supplementary Table 3). Cells were incubated with Hoescht (Thermofisher) for 1 hour at room temperature. Images were acquired using a LSM800 confocal microscope equipped with the Airyscan system (Zeiss) using a 63x objective and z-stacks. Images were analysed with Fiji and Zen.

### Electron microscopy

Cells were pre-fixed with 2% glutaraldehyde/2% PFA in Sorensen phosphate buffer 0.1M pH7.2 for 15 minutes before being fixed with 2.5% glutaraldehyde/2% PFA in Sorensen phosphate buffer 0.1M pH7.2 for 2 hours at room temperature. Cells were then washed in Sorensen phosphate buffer, resuspended in PFA 1% and sent to the METi platform (Toulouse, France). Cells were then post-fixed with 1% OsO4 in Cacodylate buffer (0.1 M, pH 7.2, EMS, Hatfield, PA), rinsed in the same buffer and pelleted, concentrated in agarose, and treated for 1 h with 2% aqueous uranyl acetate. The samples were then dehydrated in a graded ethanol series and embedded in Epon. After 48 h of polymerization at 60 °C, ultrathin sections (80 nm thick) were mounted on 200 mesh Formvar-carbon-coated copper grids. Finally, sections were stained with Uranyless and lead citrate. Grids were examined with a TEM (Jeol JEM-1400, JEOL Inc, Peabody, MA, USA) at 80 kV. Images were acquired using a digital camera (Gatan Orius, Gatan Inc, Pleasanton, CA, USA).

### ELISA

ELISA assays were performed as previously described ^43^ for the determination of the different isotype of Ig or NP-specific antibody titres in sera or culture supernatants. Briefly, plates were pre-coated with goat anti-mouse IgM (Southern Biotech) or with NP(15)-BSA (Biosearch Technologies). After a step of saturation in PBS/2%BSA, diluted sera/supernatant were added before incubation with HRP-conjugated secondary antibody. Enzymatic revelation was performed with the TMB substrate reagent set (BD OptEIA). All Antibodies used are indicated in Supplementary table 3.

### Hemagglutination inhibition (HAI) assay

Antibody titers pre and post vaccination were determined using the hemagglutination inhibition (HAI) assay using the standard WHO protocol, as previously described^44^. Sera were treated overnight with receptor-destroying enzyme (Denka Seiken Co.) and were subsequently tested by standard methods using 4 HA units of virus and a 0.5% suspension of turkey red blood cells. HAI titers were recorded as the reciprocal of the highest dilution of the serum that completely inhibited agglutination of erythrocytes by 4 HA units of the virus.

### Biomark-based transcriptomic analysis

Multiplex qPCR analysis was performed using the Biomark system (Fluidigm). Cells were sorted at 100 cells/well directly into PCR tubes containing 5μL of reverse transcription/pre-amplification mix as previously described^41^. Briefly, the mix contained 2X Reaction mix and SuperscriptIII (CellDirect One-Step qRT–PCR kit, Invitrogen) and 0,2X Taqman assay (Life technologies) (Supplementary Table 4). Targeted cDNA pre-amplification was performed for 19 cycles before processing with Dynamic Array protocol according to the manufacturer’s instructions (Fluidigm). Wells positive for *Gapdh, Actb* and control gene (*Prdm1, Irf4* and*Xpb1* for PCs and *Pax5* for B cells) expression and negative for expression of a control gene (*Cd3e*) were considered for further analysis. Mean expression of *Actb* and *Gapdh* was used for normalization. Unsupervised clustering (Spearman rank correlation test) and heatmap representation were generated with (http://www.heatmapper.ca) using the Z scores.

### RNAseq and RNAseq-based transcriptomic analysis

For RNAseq, *in vitro* generated PCs were sorted at day 2 and RNA were extracted with “RNeasy plus micro kit” (Qiagen) as recommended. RNAs were proceeded by the Integragen company for sequencing. Libraries were prepared with NEBNext® Ultra™ II Directional RNA Library Prep Kit for Illumina protocol according supplier recommendations. Briefly the key stages of this protocol were successively, the purification of PolyA containing mRNA molecules using poly-T oligo attached magnetic beads from 100ng total RNA (with the Magnetic mRNA Isolation Kit from NEB), a fragmentation using divalent cations under elevated temperature to obtain approximately 300bp pieces, double strand cDNA synthesis and finally Illumina adapters ligation and cDNA library amplification by PCR for sequencing. Sequencing was then carried out on Paired End 100b reads of Illumina NovaSeq. Image analysis and base calling was performed using Illumina Real Time Analysis (3.4.4) with default parameters.

For analysis STAR was used to obtain the number of reads associated to each gene in the Gencode vM24 annotation (restricted to protein-coding genes, antisense and lincRNAs). Raw counts for each sample were imported into R statistical software. Extracted count matrix was normalized for library size and coding length of genes to compute FPKM expression levels. The Bioconductor edgeR package was used to import raw counts into R statistical software, and compute normalized log2 CPM (counts per millions of mapped reads) using the TMM (weighted trimmed mean of M-values) as normalization procedure. The normalized expression matrix from the 500 most variant genes (based on standard deviation) was used to classify the samples according to their gene expression patterns using principal component analysis (PCA), hierarchical clustering and consensus clustering. PCA was performed by FactoMineR::PCA function with “ncp = 10, scale.unit = FALSE” parameters. Hierarchical clustering was performed by stats::hclust function (with euclidean distance and ward.D method). Differential expression analysis was performed using the Bioconductor limma package and the voom transformation. To improve the statistical power of the analysis, only genes expressed in at least one sample (FPKM >= 0.1) were considered. A q value threshold of <= 0.05 and a minimum fold change of 1.2 were used to define differentially expressed genes. Gene list from the differential analysis was ordered by decreasing log2 fold change. Gene set enrichment analysis was performed by clusterProfiler::GSEA function using the fgsea algorithm. Gene sets from MSigDB v7.2 database were selected among the C2_curated and Hallmark classes, keeping only gene sets defined by 10-500 genes.

### DropMap-based analysis of IgM secretion

Dropmap experiment were performed as described^1^, modified to detect IgM secretion from single cell. Cells from *in vitro* cultures at day2 or day 4 after LPS stimulation were centrifuged and resuspended in DropMap medium (RPMI without phenol red, supplemented with 0.1% Pluronic F68, 25 mM HEPES pH 7.4, 10% KO serum replacement (all ThermoFisher) and 0.5% recombinant human serum albumin (Sigma Aldrich). Microfluidic droplets were generated as water-in-oil emulsions using a co-flow of aqueous phases, one containing bioassay reagents (bioassay phase) and the other one containing *in vitro*-generated PCs (cell phase). *Bioassay phase*: Streptavidin-coated paramagnetic beads (300nm, Ademtech) were washed with PBS using a magnet, and then resuspended in a 1μM solution of CaptureSelect biotin anti-mouse Igk conjugate (ThermoFisher) and incubated at room temperature (RT) for 20 min. The beads were washed with PBS and resuspended in 5% pluronic F127 (ThermoFisher) and incubated for 20 min at room temperature. Following a PBS wash, the beads were resuspended in DropMap buffer and incubated at RT for 20 min. Beads were washed for the last time with PBS and resuspended in a solution of 150nM Goat anti-mouse IgM (μ chain specific) F(ab’)_2_ (Alexa647, Jackson ImmunoResearch) in DropMap buffer. *Cell phase:*To produce single-cell droplets, cell concentration was adjusted to achieve 0.3 cells per droplet in DropMap buffer. For calibration purposes with cells, purified monoclonal IgM was diluted in DropMap buffer.

Droplets were produced by hydrodynamic flow-focusing on a custom-made microfluidic device as in^1^, by co-flowing the two aqueous phases. Immediately after generation, droplets were injected into the 2D observation chamber ^1^ until filled, which was then closed for image acquisition. The droplet array in the 2D chamber was imaged using a Nikon Ti-2 Eclipse inverted microscope with motorized stage and excitation light source (Lumencor Spectra X). Fluorescence was captured using a 10x objective and a Cy5 filter, and images were recorded by a digital CMOS camera (ORCA-flash 4.0, Hamamatsu). For each time point, an array of 10×10 images was acquired, and a total of 6 acquisitions were recorded through 37.5 min. The images were analyzed with a custom Matlab script as described^1^. In brief, the ratio of fluorescent signal between the beadline and the background was estimated for every droplet at every time point. The fluorescence ratio was then used to estimate the concentration of IgM in the droplets by using a calibration curve, which was generated by measuring the fluorescent ratio from different concentrations of purified monoclonal IgM antibody.

### Statistical analysis

The p-values were determined as indicated in the figure legends using the Prism GraphPad software with the two-tailed unpaired Mann-Whitney non-parametric test for the WT vs Sec22b^B-KO^ comparison (\**p*< 0.05; **p<0.01; ***p<0.001; ****p<0.0001, “ns”= non-significant*p*-value), or with the 2way ANOVA with Sidak correction for multiple comparisons (**££**p<0.01, **£££**p<0.001).

### Supplemental Information titles and legends

**Supplementary figure 1: Raw Label-Free Quantification intensity values for the indicated proteins measured by LC-MS/MS analysis**

**Supplementary figure 2: Stx5 controls plasma cell maintenance and antibody secretion**

**Supplementary figure 3: Characterization of the *Sec22b^B-KO^* mouse model**

**Supplementary figure 4: Under-represented gene sets in Sec22b^B-KO^ PCs**

**Supplementary figure 5: Over-represented gene sets in Sec22b^B-KO^ PCs**

**Supplementary Table 1: 250 top down regulated genes in Sec22b^B-KO^ PCs**

**Supplementary table 2: 250 top up regulated genes in Sec22b^B-KO^ PCs**

**Supplementary table 3: List of antibodies used in flow cytometry, western blot and immunofluorescence**

**Supplementary table 4: List of primers used for Biomark based qPCR**

## Notes

### Competing Interest Statement

The authors have declared no competing interest.

