## supplemental data for "Plasma cell maintenance and antibody secretion are under the control of Sec22b-mediated regulation of organelle dynamics"

**Supplementary Information**

**Supplementary figure 1: Raw Label-Free Quantification intensity values for the indicated proteins measured by LC-MS/MS analysis**

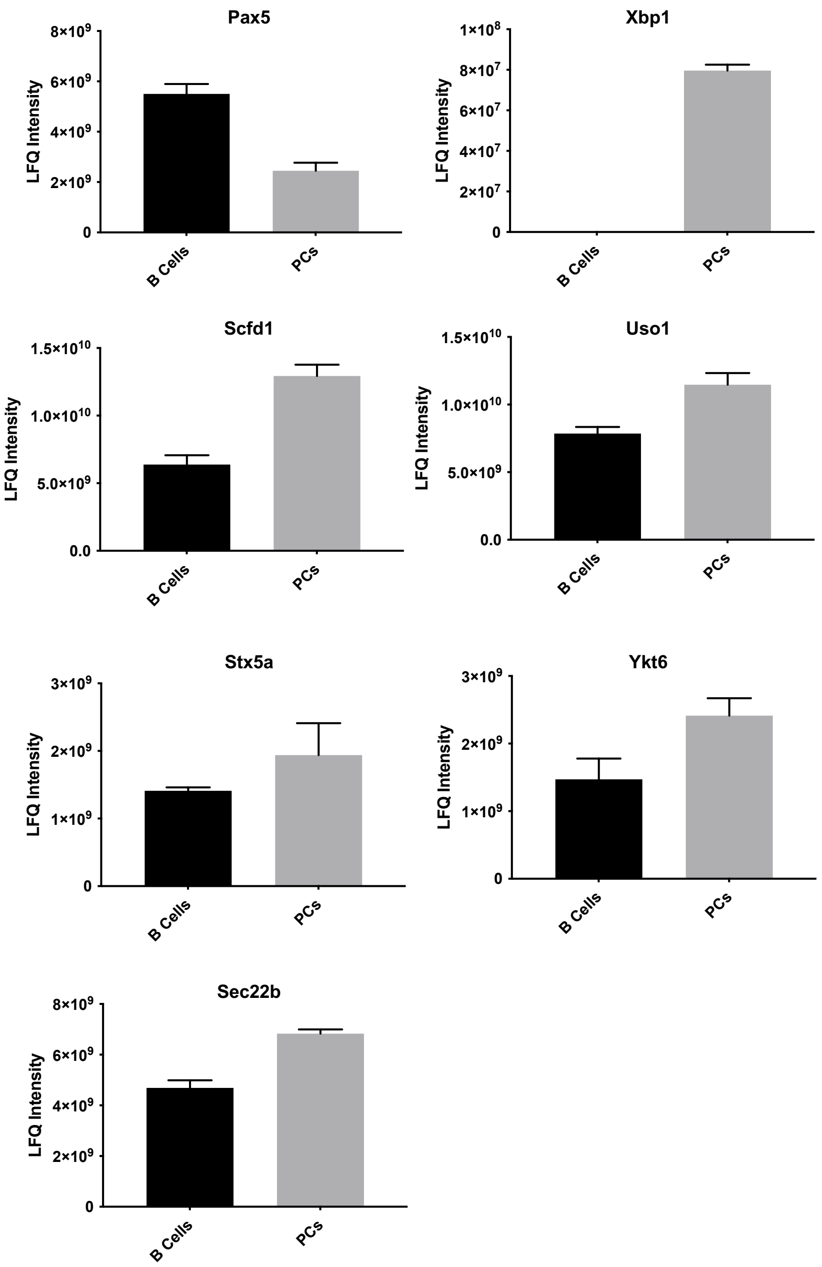

To identify proteins that exhibit altered abundance when Naive B cells differentiate into PCs, we performed label-free mass spectrometry analysis of tryptic digests of proteins isolated from splenic B cells and *in vitro* differentiated PCs (CD138 enriched). Error bars show SEM for four biological repeats.

**Supplementary figure 2: Stx5 controls plasma cell maintenance and antibody secretion**

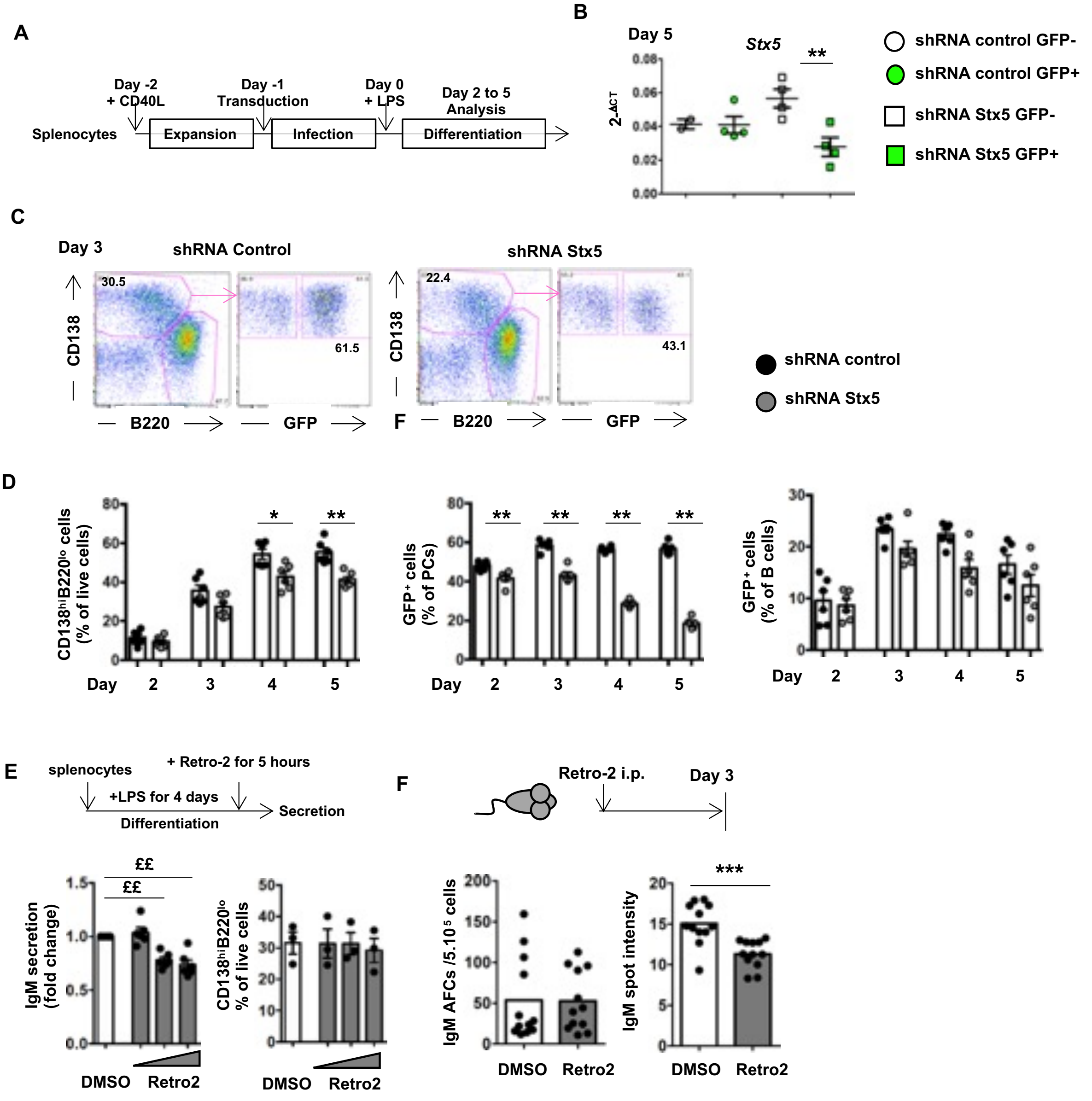

A) Schematic representation of the *in vitro* cell culture, transduction and differentiation assay.

B) Primary splenocytes were transduced with a vector encoding a control shRNA or with the vector encoding Stx5 specific shRNA together with a GFP reporter. Stx5 expression was quantified by qPCR in sorted GFP- and GFP+ cells from both experimental groups at day 5 after LPS stimulation. Data are representative of 2 independent experiments. Statistical analysis was performed with the Mann-Whitney non-parametric two-tailed unpaired test.

C-D) Representative dot plots showing the frequency of PCs (gated as $CD138^{hi}CD19^{lo}$) generated at day 3 (C) and the frequency of total PCs (D left), $GFP^{+}$ PCs (D middle) and GFP+ B cells (D right) after transduction with a control vector or with a vector encoding for a Stx5 specific shRNA at day 2, 3, 4 and 5 after transduction. N= 6 independent experiments.

E) Schematic representation of the *in vitro* cell culture and treatment with Retro-2 (top), ELISA quantification of secreted IgM in the culture supernatant (bottom left, the fold change to DMSO is shown) and frequency of PCs ($CD138^{+}B220^{low}$) determined by flow cytometry (bottom right) after 5hours of incubation with increasing doses of Retro-2 (0.3, 3 and 30µM) of *in vitro* differentiated PCs. Two independent experiments were pooled for the ELISA (n=6) and one representative experiment is shown for the flow cytometry.

F) Schematic representation of the Retro-2 *in vivo* treatment (top), quantification of the number of IgM antibody forming cells (AFCs) (bottom left) and of the intensity of IgM secretion (bottom right) in the BM of mice injected with DMSO or with Retro-2 (80mg/kg) 3 days before. Two independent experiments were pooled, n=12 mice per group. The p-values were determined with the two-tailed Mann-Whitney non-parametric test (*$p < 0.05$; **p<0.01; ***p<0.001), or with the 2way ANOVA with Sidak correction for multiple comparisons (££ p<0.01).

**Supplementary Figure 3: Characterization of the *Sec22b^B-KO^* mouse model**

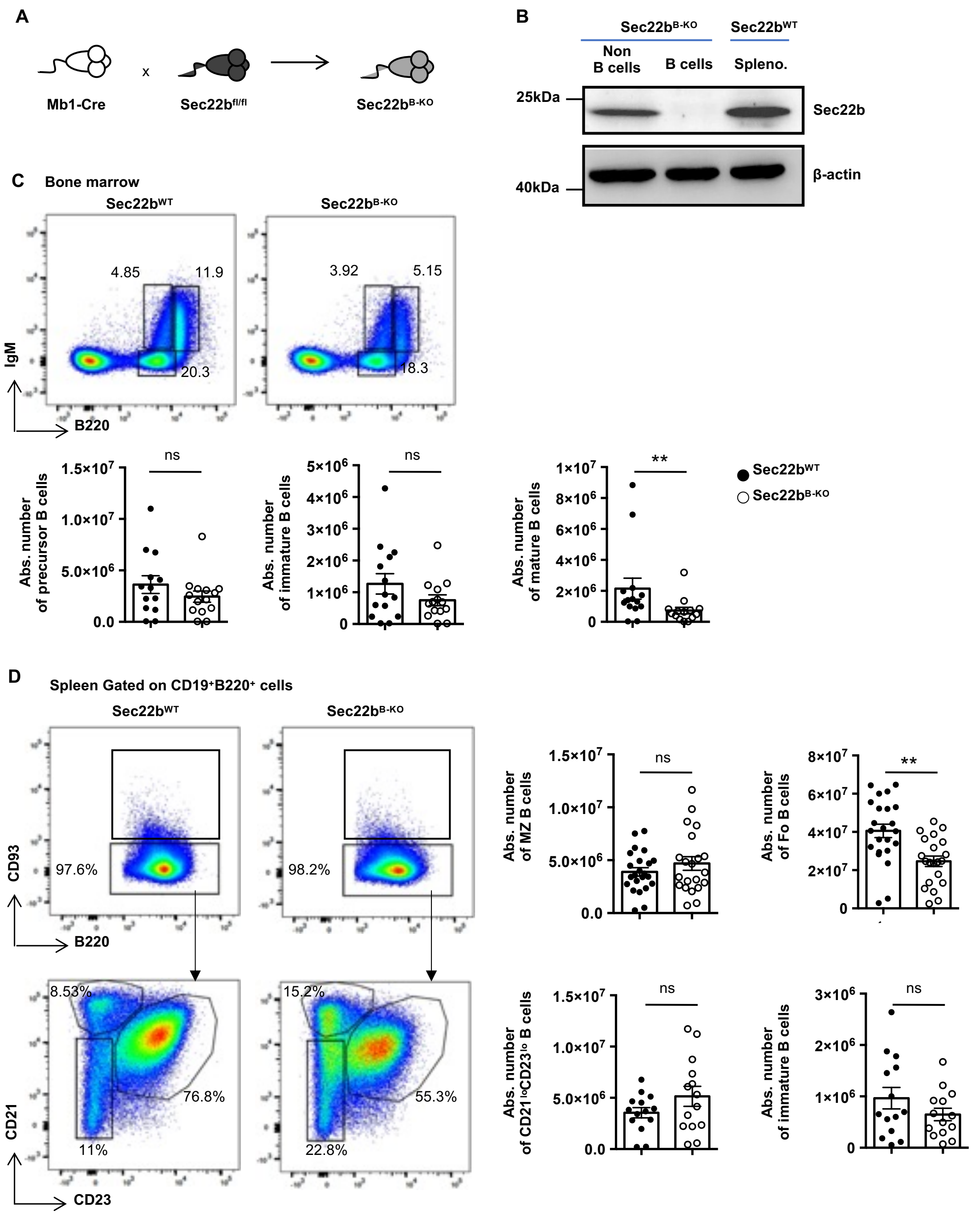

A) The Sec22b$^{B\text{-}KO}$ mouse model was generated by crossing mb1-Cre mice with Sec22b$^{fl/fl}$ mice, to obtain mice heterozygous for Cre recombinase and homozygous for Sec22b$^{fl/fl}$ (referred to as Sec22b$^{B\text{-}KO}$*)* to have a specific deletion of this SNARE protein in the B cell lineage.

B) Western blot analysis of Sec22b (top) and β-actin (bottom) from non-B cells or B cells isolated from Sec22b$^{B\text{-}KO}$ mice and total splenocytes from Sec22b$^{WT}$ mice.

C) Representative dot plots (top) and absolute number (bottom) of BM precursor B cells (B220$^{low}$IgM$^{-}$

), immature B cells (B220$^{low}$IgM$^{+}$) and mature B cells (B220$^{+}$IgM$^{+}$) in Sec22b$^{WT}$ and Sec22b$^{B\text{-}KO}$ mice.
D) Representative dot plots (left) and absolute number (right) of immature B cells (B220$^{+}$CD19$^{+}$CD93$^{+}$), follicular B cells (B220$^{+}$CD19$^{+}$CD93$^{-}$CD21$^{+}$CD23$^{+}$), marginal zone B cells (B220$^{+}$CD19$^{+}$CD93$^{-}$CD21$^{-}$CD23$^{high}$) and CD21$^{-}$CD23$^{-}$ B cells (B220$^{+}$CD19$^{+}$CD93$^{-}$CD21$^{-}$CD23$^{-}$) in Sec22b$^{WT}$ and Sec22b$^{B\text{-}KO}$ mice.

For flow cytometry experiment cells were first gated on their size and structure, their viability and doublets were excluded. n=13-23 mice from 4-5 independent experiments. The p-values were determined with the two-tailed Mann-Whitney non-parametric test. **$p < 0.01$. “ns” = non-significant *p*-value.

### Supplementary Figure 4: Under-represented gene sets in Sec22b$^{B\text{-}KO}$ PCs

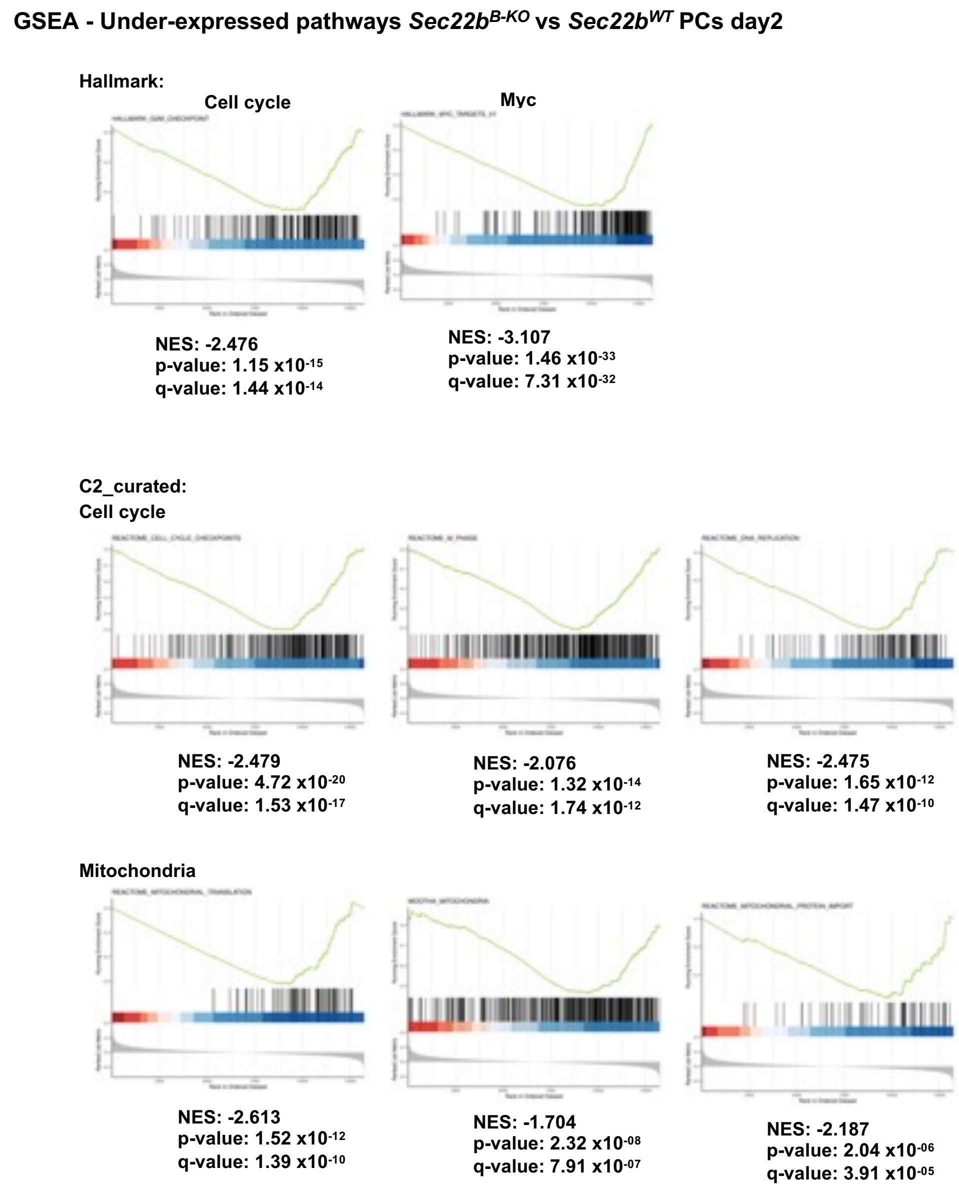

Gene set enrichment analyses were performed using the Hallmark and the C2_curated MSIgDB on RNAseq data obtained from *in vitro* generated Sec22b$^{WT}$ and Sec22b$^{B\text{-}KO}$ PCs. NES, p-values and q-values for each selected gene set are shown.

### Supplementary Figure 5: Over-represented gene sets in Sec22b$^{B\text{-}KO}$ PCs

**GSEA - Over-expressed pathways *Sec22b$^{B\text{-}KO}$* vs *Sec22b$^{WT}$* PCs day2**

**Hallmark:**

**Hallmark_protein secretion**

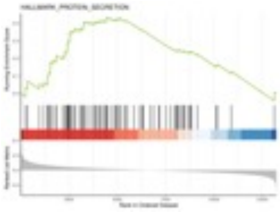

**NES: 1.596**
**p-value: 0.0021**
**q-value: 0.0096**

**C2_curated:**

**ER-Golgi transport**

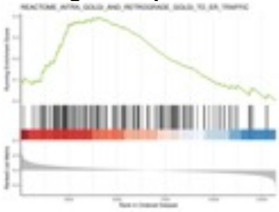

**NES: 1.608**
**p-value: 0.00046**
**q-value: 0.0044**

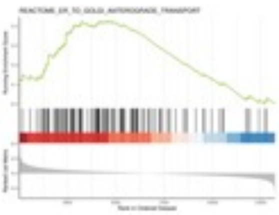

**NES: 1.662**
**p-value: 0.00054**
**q-value: 0.0050**

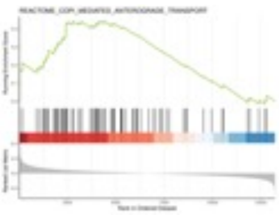

**NES: 1.625**
**p-value: 0.0034**
**q-value: 0.022**

**UPR**

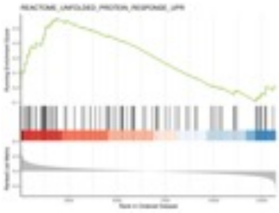

**NES: 1.740**
**p-value: 0.00035**
**q-value: 0.0035**

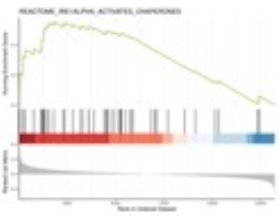

**NES: 1.868**
**p-value: 0.00054**
**q-value: 0.0051**

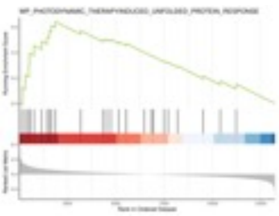

**NES: 1.900**
**p-value: 0.00084**
**q-value: 0.0072**

Gene set enrichment analyses were performed using the Hallmark and the C2_curated MSIgDB on RNAseq data obtained from *in vitro* generated Sec22b$^{WT}$ and Sec22b$^{B\text{-}KO}$ PCs. NES, p-values and q-values for each selected gene set are shown.

**Supplementary Table 1 : 250 top down regulated genes in Sec22b$^{B\text{-}KO}$ PCs**

| Gene_Name | FPKM.WT | FPKM.KO | log2FC | pval | qval |
|---|---|---|---|---|---|
| Lss | 8.56825401 | 2.87268283 | -1.4610023 | 1.2401E-11 | 2.4849E-08 |
| Hmgcs1 | 41.2146298 | 12.6431927 | -1.5902623 | 2.8482E-11 | 4.4387E-08 |
| Pcyt2 | 17.3690417 | 6.36857043 | -1.3325615 | 4.7844E-11 | 6.1005E-08 |
| Fads2 | 18.975706 | 4.07152798 | -2.1062392 | 7.3186E-11 | 7.1963E-08 |
| Cyp51 | 54.3406074 | 22.3476327 | -1.1708981 | 1.7684E-10 | 1.2148E-07 |
| Ldlr | 8.6795728 | 2.45855603 | -1.7101444 | 1.848E-10 | 1.2148E-07 |
| Msmo1 | 17.6524776 | 7.96792128 | -1.0359432 | 4.365E-10 | 1.776E-07 |
| Acat2 | 20.2603535 | 6.05876713 | -1.6241873 | 8.620E-10 | 2.4677E-07 |
| Elovl6 | 6.36330644 | 2.17111917 | -1.4369985 | 1.3888E-09 | 3.12E-07 |
| Idi1 | 21.3717499 | 6.88668479 | -1.5268591 | 1.4014E-09 | 3.12E-07 |
| Sqle | 31.146447 | 13.5704273 | -1.0905489 | 2.4411E-09 | 4.4784E-07 |
| Insig1 | 110.251597 | 47.5404861 | -1.0995955 | 3.1548E-09 | 5.2677E-07 |
| Dhcr24 | 28.7478759 | 10.3330917 | -1.37764 | 3.6204E-09 | 5.7705E-07 |
| Ldha | 393.592265 | 110.980637 | -1.7369734 | 4.7344E-09 | 6.9172E-07 |
| Stard4 | 15.137827 | 6.81370915 | -1.0386323 | 4.8045E-09 | 6.9472E-07 |
| Fgfr1op | 10.6784262 | 4.5618304 | -1.1096151 | 5.5479E-09 | 7.6996E-07 |
| Surf2 | 8.50161825 | 2.68843094 | -1.5323306 | 5.5993E-09 | 7.6996E-07 |
| Pgam1 | 65.6613951 | 22.2532633 | -1.4543746 | 5.7267E-09 | 7.7984E-07 |
| Echdc1 | 6.67061053 | 2.93570996 | -1.0688897 | 6.3202E-09 | 8.1328E-07 |
| Acsl3 | 16.7045304 | 7.92516022 | -0.9659895 | 7.6698E-09 | 9.3545E-07 |
| Apex1 | 74.6628606 | 20.6199279 | -1.7413501 | 8.681E-09 | 1.0063E-06 |
| Icam1 | 17.0459618 | 8.80207691 | -0.8387712 | 8.6538E-09 | 1.0063E-06 |
| Tspan15 | 1.49383694 | 0.56831499 | -1.2785364 | 8.6057E-09 | 1.0063E-06 |
| Dhcr7 | 10.0545011 | 5.24343935 | -0.8279307 | 9.9074E-09 | 1.0689E-06 |
| Flt3 | 5.55192425 | 2.21661128 | -1.2069847 | 1.068E-08 | 1.1263E-06 |
| Nsdhl | 12.8649607 | 5.88050199 | -1.0157331 | 1.1325E-08 | 1.1488E-06 |
| Pfkl | 25.384259 | 8.81633904 | -1.4231302 | 1.4375E-08 | 1.3441E-06 |
| Scd1 | 166.079584 | 41.6559021 | -1.915811 | 1.5057E-08 | 1.3894E-06 |
| Scarb1 | 5.03172711 | 1.96359529 | -1.2493531 | 1.5385E-08 | 1.4012E-06 |
| Ddx27 | 24.8941261 | 11.8483318 | -0.9581696 | 1.7316E-08 | 1.4719E-06 |
| Ppid | 26.6879301 | 8.13557211 | -1.6169173 | 1.8425E-08 | 1.5305E-06 |
| Cdk5r1 | 2.41313742 | 0.59678053 | -1.88986 | 1.9854E-08 | 1.6004E-06 |
| Zbtb32 | 40.5827888 | 9.75916102 | -1.9572858 | 2.0842E-08 | 1.6246E-06 |
| Fkbp3 | 12.5446757 | 5.0815353 | -1.1991922 | 2.3398E-08 | 1.7402E-06 |
| Nat10 | 14.2840458 | 5.34668556 | -1.3062485 | 2.3599E-08 | 1.7421E-06 |
| Lmo2 | 40.6125984 | 18.1293906 | -1.0471981 | 2.4732E-08 | 1.7881E-06 |
| Mybl2 | 27.223812 | 10.9252737 | -1.2175914 | 2.817E-08 | 1.9657E-06 |
| Nop16 | 40.6356111 | 16.1184395 | -1.2174837 | 2.9784E-08 | 2.0378E-06 |
| Me2 | 14.3727844 | 7.51554974 | -0.8264695 | 3.0655E-08 | 2.0771E-06 |
| Scd2 | 164.307299 | 48.1071414 | -1.6662771 | 3.1507E-08 | 2.1136E-06 |
| Fus | 25.2588217 | 11.6070257 | -1.0150247 | 3.1906E-08 | 2.1168E-06 |
| Ncl | 203.823949 | 84.5942881 | -1.1613783 | 3.3513E-08 | 2.1834E-06 |

| | | | | | |
|---|---|---|---|---|---|
| Ppp5c | 51.9729276 | 19.3188383 | -1.3238802 | 3.3693E-08 | 2.1834E-06 |
| Utp14b | 2.739302 | 1.08058333 | -1.2263403 | 3.385E-08 | 2.1834E-06 |
| Fads1 | 16.4869904 | 8.97447958 | -0.7637741 | 3.4572E-08 | 2.1855E-06 |
| Pkm | 399.846769 | 179.465117 | -1.0529468 | 3.4523E-08 | 2.1855E-06 |
| H4c9 | 38.700964 | 16.0462497 | -1.159767 | 3.5035E-08 | 2.1938E-06 |
| Prpf38b | 16.258093 | 9.46884088 | -0.666817 | 3.5443E-08 | 2.1996E-06 |
| Inf2 | 15.493966 | 7.71312821 | -0.8916687 | 3.7129E-08 | 2.2642E-06 |
| Aprt | 71.9016379 | 30.6749538 | -1.1176302 | 3.7628E-08 | 2.2847E-06 |
| Eno1 | 186.949881 | 57.476436 | -1.6227098 | 3.8733E-08 | 2.3316E-06 |
| Eno1b | 4.77195102 | 1.35046988 | -1.7096304 | 3.9072E-08 | 2.342E-06 |
| Lonp1 | 32.4179483 | 17.0426778 | -0.8191799 | 4.0741E-08 | 2.3909E-06 |
| Aacs | 13.4064779 | 7.4780055 | -0.7286582 | 4.4211E-08 | 2.4985E-06 |
| Grwd1 | 34.4037728 | 10.5583496 | -1.5913526 | 4.4252E-08 | 2.4985E-06 |
| Thop1 | 17.1790871 | 5.68553403 | -1.4923155 | 4.4494E-08 | 2.4985E-06 |
| Tpi1 | 200.078735 | 63.9924785 | -1.5531212 | 4.4806E-08 | 2.5038E-06 |
| Lipg | 1.11282125 | 0.23394749 | -2.1181924 | 4.6902E-08 | 2.5959E-06 |
| Rpp40 | 1.7478984 | 0.66677602 | -1.2753635 | 4.7439E-08 | 2.6093E-06 |
| Znhit6 | 8.92700235 | 4.11670132 | -0.9994352 | 4.7858E-08 | 2.6212E-06 |
| Wdr4 | 8.16022015 | 2.66166574 | -1.5006768 | 5.005E-08 | 2.6692E-06 |
| Cnn3 | 20.3048434 | 10.1218672 | -0.8891812 | 5.0405E-08 | 2.677E-06 |
| Ftsj3 | 42.8574198 | 15.8569984 | -1.32926 | 5.3539E-08 | 2.802E-06 |
| Bri3bp | 10.6187778 | 4.06748322 | -1.2693983 | 5.482E-08 | 2.8278E-06 |
| Mfsd2a | 13.1107067 | 4.20057975 | -1.5476844 | 5.5798E-08 | 2.8582E-06 |
| Pdxk | 4.49105218 | 1.47523748 | -1.4961434 | 5.6151E-08 | 2.8639E-06 |
| Rpf2 | 33.085641 | 11.9939701 | -1.3517842 | 5.7114E-08 | 2.8746E-06 |
| Pa2g4 | 157.989551 | 64.8145755 | -1.1815235 | 5.8494E-08 | 2.9197E-06 |
| Dctpp1 | 104.449569 | 41.5536628 | -1.2216924 | 5.8752E-08 | 2.9222E-06 |
| Sms | 9.92595213 | 4.63665307 | -0.9770364 | 5.967E-08 | 2.9573E-06 |
| Mrpl42 | 19.0662535 | 8.79094583 | -1.0058073 | 6.02E-08 | 2.9731E-06 |
| Ranbp1 | 69.1663027 | 28.1863745 | -1.1963613 | 6.1693E-08 | 3.0362E-06 |
| Marcksl1 | 67.6115029 | 16.5464035 | -1.9421071 | 6.2402E-08 | 3.0424E-06 |
| Mif | 444.464436 | 180.882376 | -1.1984281 | 6.3179E-08 | 3.0663E-06 |
| Gmip | 8.99818043 | 4.87243959 | -0.7683209 | 6.5782E-08 | 3.1598E-06 |
| Rrp1b | 12.4511431 | 3.98094015 | -1.5597112 | 6.6686E-08 | 3.1745E-06 |
| Slc20a1 | 8.91007222 | 4.51009675 | -0.868933 | 6.7554E-08 | 3.2011E-06 |
| Tkt | 74.2154403 | 36.1727783 | -0.9279239 | 7.1054E-08 | 3.3112E-06 |
| Ydjc | 2.45014308 | 0.75441484 | -1.567073 | 7.1976E-08 | 3.325E-06 |
| Gypc | 24.3544816 | 5.114789 | -2.1513416 | 7.4045E-08 | 3.4051E-06 |
| Pmvk | 23.4429288 | 12.4607926 | -0.7994702 | 7.4342E-08 | 3.4076E-06 |
| Traf4 | 18.0326208 | 8.87540096 | -0.9110618 | 7.5373E-08 | 3.4341E-06 |
| Shmt1 | 30.9301267 | 7.21985001 | -2.018633 | 7.6658E-08 | 3.4684E-06 |
| Wdr12 | 6.10414455 | 2.55028156 | -1.1411704 | 7.7098E-08 | 3.4771E-06 |
| Farsa | 21.1588978 | 11.4584934 | -0.7714129 | 7.8651E-08 | 3.5021E-06 |
| Snrnp70 | 40.9377394 | 21.0592483 | -0.8508479 | 8.0516E-08 | 3.5738E-06 |
| 1110038B12Rik | 28.3962719 | 13.6917727 | -0.9373481 | 8.2723E-08 | 3.6602E-06 |

| | | | | | |
|---|---|---|---|---|---|
| Nbeal2 | 2.84961182 | 1.2279435 | -1.0983979 | 8.4425E-08 | 3.7005E-06 |
| Isyna1 | 75.4849984 | 30.5360452 | -1.2085092 | 8.5468E-08 | 3.7315E-06 |
| Nop56 | 69.6960607 | 25.0986184 | -1.3767177 | 8.5665E-08 | 3.7315E-06 |
| Epcam | 13.5267062 | 6.45817477 | -0.9461685 | 8.6845E-08 | 3.7396E-06 |
| Gm17494 | 0.81248176 | 0.21012953 | -1.8300529 | 8.9296E-08 | 3.7717E-06 |
| Mtap | 37.2196081 | 15.4490726 | -1.1580458 | 8.9547E-08 | 3.7717E-06 |
| Nsun5 | 6.28065341 | 2.78564997 | -1.0540886 | 9.3574E-08 | 3.8946E-06 |
| Pum3 | 14.4612984 | 6.43405529 | -1.0614411 | 9.4464E-08 | 3.92E-06 |
| Pgs1 | 9.90426581 | 5.79722283 | -0.6578622 | 9.565E-08 | 3.9575E-06 |
| Cacybp | 82.1216986 | 30.058963 | -1.3420252 | 1.0206E-07 | 4.1306E-06 |
| Mvd | 25.0809168 | 13.5491604 | -0.7781807 | 1.0117E-07 | 4.1306E-06 |
| Pus7 | 7.8476253 | 2.68193855 | -1.4377053 | 1.0219E-07 | 4.1306E-06 |
| Nudc | 152.269089 | 63.8001613 | -1.1534943 | 1.0279E-07 | 4.1428E-06 |
| Gapdh | 461.304721 | 240.23328 | -0.8326048 | 1.0488E-07 | 4.1723E-06 |
| Hmgn1 | 93.4987252 | 51.0894805 | -0.7611395 | 1.048E-07 | 4.1723E-06 |
| Sc5d | 12.0999582 | 6.95568025 | -0.684767 | 1.0656E-07 | 4.1984E-06 |
| Ddx11 | 4.29229073 | 2.12125418 | -0.8978846 | 1.1414E-07 | 4.4396E-06 |
| Pnn | 23.0757921 | 11.9046688 | -0.846196 | 1.1442E-07 | 4.4396E-06 |
| Crtc2 | 16.9579125 | 9.89598325 | -0.6626003 | 1.1598E-07 | 4.4609E-06 |
| Gart | 57.972664 | 20.6816937 | -1.3859317 | 1.1991E-07 | 4.558E-06 |
| Atad3a | 28.1887967 | 10.1383583 | -1.3744741 | 1.2064E-07 | 4.5731E-06 |
| Hsp90aa1 | 255.105755 | 89.7493845 | -1.4181893 | 1.2211E-07 | 4.5919E-06 |
| Snhg12 | 8.44866987 | 4.22167947 | -0.8833022 | 1.2376E-07 | 4.6291E-06 |
| Tagln2 | 139.55187 | 73.8092625 | -0.8071984 | 1.2375E-07 | 4.6291E-06 |
| Serbp1 | 41.4719696 | 20.2192668 | -0.9270339 | 1.2535E-07 | 4.6519E-06 |
| Slc29a1 | 10.8451604 | 4.61380416 | -1.1208001 | 1.2534E-07 | 4.6519E-06 |
| Wdr77 | 6.82500732 | 2.93431452 | -1.1041624 | 1.2537E-07 | 4.6519E-06 |
| Nsun2 | 58.2075831 | 23.40093 | -1.2121876 | 1.3414E-07 | 4.938E-06 |
| Mrto4 | 39.5360771 | 14.1046177 | -1.3842843 | 1.3775E-07 | 5.0448E-06 |
| Noc2l | 54.9085113 | 22.3461613 | -1.1979513 | 1.3751E-07 | 5.0448E-06 |
| Cct8 | 156.429099 | 70.8344859 | -1.0427393 | 1.394E-07 | 5.0637E-06 |
| Fubp1 | 14.5763978 | 6.31556817 | -1.101349 | 1.3946E-07 | 5.0637E-06 |
| Pwp2 | 17.403609 | 7.11706501 | -1.1798677 | 1.3953E-07 | 5.0637E-06 |
| Carm1 | 21.1767643 | 10.6469565 | -0.8860249 | 1.4498E-07 | 5.0759E-06 |
| Dctd | 5.06466071 | 1.30827224 | -1.8220852 | 1.4324E-07 | 5.0759E-06 |
| Fcer2a | 4.94588061 | 0.73456418 | -2.6976286 | 1.4348E-07 | 5.0759E-06 |
| Gar1 | 41.9545035 | 17.6328305 | -1.1457304 | 1.4474E-07 | 5.0759E-06 |
| Gpn1 | 4.48627678 | 2.17971584 | -0.9274573 | 1.4197E-07 | 5.0759E-06 |
| Hsd17b7 | 5.32479599 | 1.89669135 | -1.3976029 | 1.4406E-07 | 5.0759E-06 |
| Pdcd5 | 19.7635177 | 11.0586376 | -0.7230124 | 1.438E-07 | 5.0759E-06 |
| Sema7a | 25.2682779 | 12.7709944 | -0.8749612 | 1.4858E-07 | 5.1583E-06 |
| Ak2 | 87.5968588 | 41.6964534 | -0.9638116 | 1.5025E-07 | 5.1905E-06 |
| Dbi | 63.3523148 | 31.5611039 | -0.8936292 | 1.5424E-07 | 5.2786E-06 |
| Fgd2 | 8.96809311 | 2.80076271 | -1.5533949 | 1.5485E-07 | 5.2786E-06 |
| Hspd1 | 280.824307 | 97.0305934 | -1.4529475 | 1.5505E-07 | 5.2786E-06 |

| | | | | | |
|---|---|---|---|---|---|
| Prag1 | 7.21418876 | 3.43495381 | -0.9593574 | 1.5391E-07 | 5.2786E-06 |
| Hirip3 | 31.7030198 | 14.7276123 | -1.0076799 | 1.605E-07 | 5.3108E-06 |
| Nasp | 34.3617743 | 14.1421082 | -1.1874818 | 1.5762E-07 | 5.3108E-06 |
| Trap1 | 54.8779605 | 28.3681383 | -0.8416449 | 1.6119E-07 | 5.3108E-06 |
| Utp4 | 29.4714393 | 10.7522815 | -1.3504047 | 1.5794E-07 | 5.3108E-06 |
| 3110082I17Rik | 9.6509888 | 3.27484572 | -1.4453386 | 1.6219E-07 | 5.3276E-06 |
| Mat2a | 51.7360175 | 19.6367395 | -1.2907867 | 1.6639E-07 | 5.3772E-06 |
| Wdr46 | 30.1119942 | 13.6781825 | -1.0309348 | 1.6678E-07 | 5.3776E-06 |
| Anp32a | 73.6579402 | 33.7832778 | -1.0216852 | 1.6783E-07 | 5.3872E-06 |
| Grap | 66.0028612 | 34.2189432 | -0.8293041 | 1.7144E-07 | 5.4481E-06 |
| Slc2a1 | 17.5323248 | 9.85000774 | -0.7156347 | 1.7767E-07 | 5.6125E-06 |
| Acy1 | 6.40415738 | 2.25328174 | -1.4020424 | 1.7909E-07 | 5.6226E-06 |
| Slc29a2 | 1.07433067 | 0.17848101 | -2.4613394 | 1.7919E-07 | 5.6226E-06 |
| Snhg3 | 13.3713253 | 3.80653698 | -1.7136906 | 1.8027E-07 | 5.644E-06 |
| A630001G21Rik | 27.5241587 | 15.6243202 | -0.701754 | 1.812E-07 | 5.6575E-06 |
| Gemin6 | 27.4336311 | 13.3229124 | -0.9302028 | 1.8151E-07 | 5.6575E-06 |
| Pnpt1 | 20.4644811 | 8.72513327 | -1.1207023 | 1.8715E-07 | 5.7693E-06 |
| Fam111a | 34.7621613 | 14.7941072 | -1.1324791 | 1.8855E-07 | 5.7994E-06 |
| Snrpa1 | 24.9647853 | 11.6509704 | -0.9926056 | 1.9069E-07 | 5.8397E-06 |
| Pgd | 20.9643447 | 9.9754339 | -0.9566437 | 1.9122E-07 | 5.8432E-06 |
| Acaca | 5.08754614 | 2.45588342 | -0.9447636 | 1.9337E-07 | 5.896E-06 |
| Fbl | 37.5159634 | 14.8553059 | -1.2331585 | 1.9535E-07 | 5.9308E-06 |
| Set | 174.478943 | 75.389788 | -1.1075613 | 1.9672E-07 | 5.9465E-06 |
| Fdft1 | 15.6190544 | 8.82099841 | -0.7132296 | 1.9978E-07 | 6.0002E-06 |
| Ppan | 25.1963056 | 12.5514582 | -0.8992079 | 2.1015E-07 | 6.2316E-06 |
| Exosc1 | 11.2718921 | 5.32594244 | -0.9673966 | 2.1384E-07 | 6.3131E-06 |
| Lyar | 68.499119 | 27.0162705 | -1.2414494 | 2.215E-07 | 6.4229E-06 |
| Sephs2 | 58.3709451 | 32.0605618 | -0.7497708 | 2.2074E-07 | 6.4229E-06 |
| Ddx55 | 4.69351957 | 2.4646651 | -0.8150927 | 2.2668E-07 | 6.5286E-06 |
| Dnajc2 | 29.2803674 | 14.9192668 | -0.8673129 | 2.2968E-07 | 6.5478E-06 |
| Dph2 | 7.05286765 | 3.63434931 | -0.8418329 | 2.2913E-07 | 6.5478E-06 |
| Nop2 | 52.5885297 | 20.3055697 | -1.2622746 | 2.2884E-07 | 6.5478E-06 |
| Uhrf1 | 75.7194421 | 35.5059418 | -0.994123 | 2.2968E-07 | 6.5478E-06 |
| Cluh | 25.3568144 | 8.58660034 | -1.4756737 | 2.351E-07 | 6.6887E-06 |
| Ddx51 | 8.47567661 | 3.69402117 | -1.0739422 | 2.3558E-07 | 6.6888E-06 |
| Dkc1 | 39.8639743 | 14.4054823 | -1.3735196 | 2.3687E-07 | 6.7118E-06 |
| Cyth4 | 15.4093794 | 8.37167125 | -0.7706545 | 2.3972E-07 | 6.7533E-06 |
| Tfdp1 | 78.4400917 | 35.250557 | -1.0554852 | 2.4103E-07 | 6.7613E-06 |
| Prpf3 | 11.8770045 | 6.31080232 | -0.8030732 | 2.4451E-07 | 6.8453E-06 |
| Rrp12 | 19.9857853 | 7.5280107 | -1.3074798 | 2.5147E-07 | 6.9568E-06 |
| Setd6 | 7.59376952 | 2.63229132 | -1.4140686 | 2.5144E-07 | 6.9568E-06 |
| U2af1 | 52.6335964 | 27.1078286 | -0.8538408 | 2.5132E-07 | 6.9568E-06 |
| Ddx21 | 101.707977 | 45.9277155 | -1.0458448 | 2.5357E-07 | 6.9653E-06 |
| Dph5 | 3.27185564 | 1.51218776 | -0.9893894 | 2.5426E-07 | 6.9653E-06 |
| Ripk3 | 27.5796962 | 15.4859769 | -0.7212982 | 2.5245E-07 | 6.9653E-06 |

| | | | | | |
|---|---|---|---|---|---|
| Bzw2 | 24.2934115 | 10.0132953 | -1.1776276 | 2.6184E-07 | 7.0683E-06 |
| Gpatch4 | 24.1588158 | 10.0074096 | -1.1673468 | 2.6191E-07 | 7.0683E-06 |
| Polr3g | 3.59767821 | 1.47671818 | -1.1588163 | 2.6199E-07 | 7.0683E-06 |
| Rsl1d1 | 104.282222 | 45.7529902 | -1.0823063 | 2.6023E-07 | 7.0683E-06 |
| Polr1b | 13.9586963 | 5.32160318 | -1.2849283 | 2.661E-07 | 7.1227E-06 |
| Wdr18 | 16.6156591 | 8.48377892 | -0.861252 | 2.6561E-07 | 7.1227E-06 |
| Atic | 27.831701 | 9.63385869 | -1.4437515 | 2.6841E-07 | 7.1376E-06 |
| Nhp2 | 133.783012 | 50.1121426 | -1.3038279 | 2.8276E-07 | 7.3991E-06 |
| Cd3eap | 11.0155385 | 3.73219713 | -1.4547488 | 2.8961E-07 | 7.4945E-06 |
| Exosc2 | 22.9074875 | 9.79700992 | -1.1144129 | 2.9198E-07 | 7.4988E-06 |
| Shisa8 | 28.7109486 | 10.2772456 | -1.3666173 | 2.9136E-07 | 7.4988E-06 |
| Acly | 62.6426972 | 36.7993259 | -0.6581826 | 2.9694E-07 | 7.5622E-06 |
| Chaf1b | 9.77405337 | 4.40342274 | -1.0399261 | 2.9872E-07 | 7.5622E-06 |
| Phb2 | 86.2160716 | 42.4177006 | -0.9084764 | 2.9679E-07 | 7.5622E-06 |
| Tra2a | 16.3512252 | 9.13556906 | -0.7289314 | 2.985E-07 | 7.5622E-06 |
| Ybx3 | 150.447731 | 64.2561079 | -1.1313189 | 2.9923E-07 | 7.5622E-06 |
| Snhg1 | 28.2846716 | 14.5070464 | -0.8515614 | 3.0426E-07 | 7.6479E-06 |
| Bop1 | 60.4070115 | 27.5585146 | -1.024434 | 3.0643E-07 | 7.677E-06 |
| Noc4l | 32.7978959 | 13.7458394 | -1.1449604 | 3.076E-07 | 7.677E-06 |
| Tnfrsf22 | 1.66451967 | 0.63650961 | -1.2771072 | 3.1222E-07 | 7.7371E-06 |
| Tomm40 | 44.066503 | 19.233421 | -1.0999855 | 3.1125E-07 | 7.7371E-06 |
| Lactb2 | 6.02471241 | 2.92714593 | -0.9238601 | 3.1585E-07 | 7.7739E-06 |
| Sp110 | 18.2486258 | 9.12240564 | -0.8904227 | 3.1514E-07 | 7.7739E-06 |
| Ptpn6 | 61.2230187 | 25.0689157 | -1.1788882 | 3.1875E-07 | 7.816E-06 |
| Tardbp | 43.2135414 | 22.1910373 | -0.8582379 | 3.2934E-07 | 8.0065E-06 |
| Uchl5 | 18.5742391 | 9.15139617 | -0.90832 | 3.2937E-07 | 8.0065E-06 |
| Nop10 | 70.8135977 | 29.4845157 | -1.1487389 | 3.3141E-07 | 8.0134E-06 |
| Pfas | 14.9393854 | 6.29193403 | -1.1521317 | 3.3157E-07 | 8.0134E-06 |
| Rnps1 | 36.0336989 | 19.7844529 | -0.7580303 | 3.3085E-07 | 8.0134E-06 |
| Ddx39b | 65.8868456 | 37.1617358 | -0.7183831 | 3.3411E-07 | 8.0243E-06 |
| Cdca7 | 28.7154846 | 8.86420421 | -1.593075 | 3.3471E-07 | 8.0248E-06 |
| Ppat | 25.0878496 | 9.80951587 | -1.2613522 | 3.3681E-07 | 8.0478E-06 |
| Akap8 | 14.3351994 | 7.69481542 | -0.786104 | 3.4231E-07 | 8.0631E-06 |
| Dek | 58.9413279 | 23.8490641 | -1.1989629 | 3.4262E-07 | 8.0631E-06 |
| Dhodh | 4.0529472 | 1.91596783 | -0.9649886 | 3.4012E-07 | 8.0631E-06 |
| Otud6b | 21.9949089 | 12.240962 | -0.734482 | 3.3863E-07 | 8.0631E-06 |
| Pdcd11 | 15.1730927 | 8.20718644 | -0.7782819 | 3.3983E-07 | 8.0631E-06 |
| Mybbp1a | 112.341809 | 45.3315558 | -1.2075281 | 3.4527E-07 | 8.1105E-06 |
| Rbm19 | 12.3681261 | 5.75482361 | -0.9998359 | 3.4579E-07 | 8.1105E-06 |
| Lsm7 | 12.7719719 | 5.22323763 | -1.1710838 | 3.4747E-07 | 8.1232E-06 |
| Pif1 | 7.94548809 | 3.99930293 | -0.8791316 | 3.4749E-07 | 8.1232E-06 |
| Arglu1 | 15.6442207 | 7.37871689 | -0.9687233 | 3.4907E-07 | 8.1464E-06 |
| Mettl16 | 5.78411786 | 2.57338555 | -1.060601 | 3.5388E-07 | 8.2328E-06 |
| Slc6a6 | 8.21250997 | 4.53053398 | -0.7436557 | 3.5588E-07 | 8.2497E-06 |
| Igsf8 | 5.26673181 | 2.53394636 | -0.9429703 | 3.5815E-07 | 8.2758E-06 |

| | | | | | |
|---|---|---|---|---|---|
| Elac2 | 18.6556986 | 8.76591846 | -0.9795827 | 3.6E-07 | 8.2763E-06 |
| Npm1 | 641.680216 | 321.760204 | -0.8850113 | 3.5959E-07 | 8.2763E-06 |
| Rrp9 | 26.9895929 | 9.18592957 | -1.4494495 | 3.5996E-07 | 8.2763E-06 |
| Zfp608 | 1.71133655 | 0.82776843 | -0.927823 | 3.6053E-07 | 8.2763E-06 |
| Luc7l3 | 16.6864888 | 9.901422 | -0.637643 | 3.6113E-07 | 8.2764E-06 |
| Rrs1 | 42.0211858 | 18.4623088 | -1.0771518 | 3.6311E-07 | 8.2947E-06 |
| Tars2 | 8.74745541 | 4.93144657 | -0.7171052 | 3.6254E-07 | 8.2947E-06 |
| C1qbp | 120.598641 | 50.2476547 | -1.1640199 | 3.6629E-07 | 8.3268E-06 |
| Nip7 | 24.5138904 | 11.0160419 | -1.0354401 | 3.6711E-07 | 8.3319E-06 |
| Tbrg4 | 36.8265773 | 18.7357568 | -0.8677646 | 3.7077E-07 | 8.4013E-06 |
| Dis3 | 13.3028362 | 5.80618046 | -1.0952203 | 3.7159E-07 | 8.4062E-06 |
| Lbhd1 | 3.90550219 | 1.87189609 | -0.9453705 | 3.7824E-07 | 8.5292E-06 |
| Agpat5 | 5.47982335 | 2.86237079 | -0.8275443 | 3.8918E-07 | 8.72E-06 |
| Fabp5 | 5.70244802 | 1.61362848 | -1.7075375 | 3.8994E-07 | 8.723E-06 |
| Amd1 | 23.1959124 | 10.989617 | -0.9728992 | 3.9643E-07 | 8.8119E-06 |
| Stip1 | 170.703396 | 75.2983801 | -1.078698 | 4.0421E-07 | 8.9423E-06 |
| Cd40 | 10.4896306 | 3.49203991 | -1.4361713 | 4.0538E-07 | 8.954E-06 |
| Cct6a | 133.369771 | 58.8690206 | -1.0694067 | 4.1025E-07 | 8.973E-06 |
| Ebna1bp2 | 31.9362662 | 14.6199572 | -1.0213358 | 4.1136E-07 | 8.973E-06 |
| Eef1aknmt | 3.8005288 | 1.74309112 | -1.0168398 | 4.0951E-07 | 8.973E-06 |
| Nop58 | 29.6806354 | 13.4691936 | -1.0373355 | 4.1135E-07 | 8.973E-06 |
| Tuba1b | 207.392068 | 85.0044206 | -1.1849099 | 4.1332E-07 | 9.0018E-06 |
| Impdh2 | 90.7190753 | 46.4679314 | -0.8541398 | 4.1653E-07 | 9.0437E-06 |
| Nap1l1 | 113.513733 | 62.0763688 | -0.7620347 | 4.193E-07 | 9.0758E-06 |
| Kat2a | 18.5453048 | 8.85440777 | -0.9626712 | 4.2071E-07 | 9.0922E-06 |
| 2810004N23Rik | 9.68216348 | 4.88400704 | -0.8690086 | 4.2295E-07 | 9.0986E-06 |
| Acsl4 | 17.1770638 | 10.8478202 | -0.549714 | 4.2256E-07 | 9.0986E-06 |
| Dnaja1 | 16.5014827 | 8.08866449 | -0.9139051 | 4.2263E-07 | 9.0986E-06 |

**Supplementary Table 2 : 250 top up regulated genes in Sec22b$^{B\text{-}KO}$ PCs**

| GEnE_NamE | FPKM.WT | FPKM.KO | log2FC | pval | qval |
|---|---|---|---|---|---|
| Rcn3 | 3.95244129 | 28.2256264 | 2.94541415 | 3.6538E-13 | 5.1248E-09 |
| Aldh1l2 | 0.97068046 | 14.9693242 | 4.06771228 | 9.7284E-13 | 6.8225E-09 |
| Anxa5 | 19.6312692 | 67.6798591 | 1.90267766 | 5.8389E-12 | 1.6379E-08 |
| Ighv1-12 | 91.3141777 | 262.873354 | 1.63770725 | 3.6498E-12 | 1.6379E-08 |
| KdElr3 | 2.89563443 | 32.0034861 | 3.58792461 | 5.6699E-12 | 1.6379E-08 |
| Hapln4 | 0.35185669 | 3.1586156 | 3.26100562 | 7.1344E-12 | 1.6678E-08 |
| Atf5 | 37.1824171 | 186.050449 | 2.43096651 | 2.3181E-11 | 4.0642E-08 |
| Ighv1-74 | 158.191774 | 407.725183 | 1.48005927 | 4.2968E-11 | 6.0267E-08 |
| Pck2 | 30.2316111 | 123.439586 | 2.13980755 | 6.8118E-11 | 7.1963E-08 |
| Ptov1 | 0.83255733 | 5.15312938 | 2.72882675 | 6.7398E-11 | 7.1963E-08 |
| VEgfa | 16.4845734 | 44.916135 | 1.56414288 | 7.696E-11 | 7.1963E-08 |
| Fcrla | 30.0184528 | 70.5261476 | 1.34403108 | 1.1394E-10 | 9.7678E-08 |
| Pycr1 | 13.500785 | 51.9078507 | 2.06442437 | 1.2535E-10 | 9.7678E-08 |
| Tmc6 | 6.75469967 | 14.7802354 | 1.24056449 | 1.2089E-10 | 9.7678E-08 |
| Hdac9 | 2.09836795 | 7.90746978 | 2.01222542 | 1.7737E-10 | 1.2148E-07 |
| Ighv1-78 | 175.033675 | 523.163136 | 1.69867469 | 1.9054E-10 | 1.2148E-07 |
| Dusp5 | 7.48036711 | 16.5835982 | 1.26197664 | 2.0654E-10 | 1.2596E-07 |
| Rassf4 | 3.35563798 | 7.90633389 | 1.34615616 | 2.4681E-10 | 1.4424E-07 |
| SElEnom | 5.10934057 | 17.8335482 | 1.91632104 | 3.0367E-10 | 1.7037E-07 |
| MagEd2 | 0.55352151 | 6.23833266 | 3.57222716 | 3.2248E-10 | 1.7397E-07 |
| B3gnt9 | 5.26744261 | 21.6136886 | 2.14346305 | 4.5583E-10 | 1.776E-07 |
| Cd28 | 1.30171125 | 7.0354162 | 2.53282616 | 3.6945E-10 | 1.776E-07 |
| CEbpb | 24.8524107 | 61.4684689 | 1.41125685 | 3.8996E-10 | 1.776E-07 |
| Ighv8-8 | 296.018709 | 1102.1792 | 1.99658704 | 3.4409E-10 | 1.776E-07 |
| MagEd1 | 7.11806636 | 24.2847606 | 1.86880346 | 4.4644E-10 | 1.776E-07 |
| PlEkha1 | 2.15696285 | 5.23836059 | 1.39050571 | 4.1291E-10 | 1.776E-07 |
| Plxna1 | 4.27809191 | 14.4071916 | 1.86793714 | 3.9552E-10 | 1.776E-07 |
| Rrm2b | 2.58130441 | 6.10816923 | 1.35127282 | 4.1430E-10 | 1.776E-07 |
| Sft2d2 | 34.4282347 | 94.6883489 | 1.57779801 | 4.5466E-10 | 1.776E-07 |
| Fam129a | 4.81711927 | 18.273281 | 2.02779748 | 4.7637E-10 | 1.8059E-07 |
| Fam114a1 | 0.76205666 | 5.12239184 | 2.85050222 | 5.3447E-10 | 1.9704E-07 |
| Ighv3-8 | 77.7134879 | 326.521966 | 2.20463601 | 5.6191E-10 | 1.9704E-07 |
| SEsn2 | 14.7650456 | 42.4472139 | 1.63728825 | 5.4822E-10 | 1.9704E-07 |
| Evi2a | 18.0110335 | 60.8388687 | 1.85330884 | 5.7990E-10 | 1.9839E-07 |
| SEc23a | 17.8776281 | 41.4506525 | 1.32789755 | 6.2665E-10 | 2.0297E-07 |
| Tifa | 2.09109694 | 5.69958631 | 1.55636589 | 6.2694E-10 | 2.0297E-07 |
| Zfp9 | 1.32972835 | 4.3112639 | 1.79831354 | 6.3671E-10 | 2.0297E-07 |
| Copz2 | 0.47058406 | 5.95454045 | 3.74816703 | 6.7108E-10 | 2.0462E-07 |
| Ighv1-58 | 47.8972969 | 152.838548 | 1.77586681 | 6.6684E-10 | 2.0462E-07 |
| Lasp1 | 6.64790742 | 11.9180495 | 0.95523756 | 6.8654E-10 | 2.0488E-07 |
| Ikbip | 1.87484725 | 4.98237371 | 1.52239466 | 8.4695E-10 | 2.4677E-07 |
| CtsE | 3.51488042 | 9.15376791 | 1.49343696 | 9.1224E-10 | 2.559E-07 |
| Igkv12-98 | 107.721563 | 263.505214 | 1.40215105 | 9.3287E-10 | 2.5656E-07 |

| | | | | | |
|---|---|---|---|---|---|
| Abr | 2.51223188 | 5.24500495 | 1.17245331 | 1.0646E-09 | 2.6198E-07 |
| BhlhE41 | 4.74207339 | 10.1582443 | 1.20860153 | 1.053E-09 | 2.6198E-07 |
| Irgq | 8.82528071 | 21.3618862 | 1.39116172 | 1.0566E-09 | 2.6198E-07 |
| Slc3a2 | 111.000524 | 280.504692 | 1.44873381 | 1.0405E-09 | 2.6198E-07 |
| Slc6a9 | 1.9816791 | 10.5969137 | 2.51914349 | 9.9537E-10 | 2.6198E-07 |
| TmEm263 | 15.8314176 | 30.2251103 | 1.04471015 | 1.0584E-09 | 2.6198E-07 |
| Chpf2 | 15.8208273 | 32.512173 | 1.15506603 | 1.0876E-09 | 2.6301E-07 |
| Faah | 3.57928509 | 7.79686525 | 1.23410039 | 1.1463E-09 | 2.725E-07 |
| ArfgEf3 | 1.18740969 | 6.65536999 | 2.62340302 | 1.2781E-09 | 2.9879E-07 |
| Dap | 43.035225 | 111.589761 | 1.49780095 | 1.3962E-09 | 3.12E-07 |
| MlEc | 88.0293766 | 149.8739 | 0.88147187 | 1.5084E-09 | 3.2548E-07 |
| Mxd4 | 8.35434174 | 25.9818553 | 1.7554385 | 1.4977E-09 | 3.2548E-07 |
| Igkv16-104 | 680.272709 | 1459.30687 | 1.21156747 | 1.6217E-09 | 3.4463E-07 |
| Chac1 | 29.129383 | 82.906231 | 1.62728404 | 1.6769E-09 | 3.4588E-07 |
| S100a11 | 55.1503858 | 151.389057 | 1.57104305 | 1.6753E-09 | 3.4588E-07 |
| Igkv4-50 | 141.049356 | 449.891241 | 1.77035923 | 1.7417E-09 | 3.5129E-07 |
| TmEm26 | 1.1636833 | 3.8498455 | 1.84833775 | 1.7532E-09 | 3.5129E-07 |
| Slc7a11 | 0.61297132 | 3.0328239 | 2.42421242 | 1.807E-09 | 3.5698E-07 |
| LEprotl1 | 14.9103053 | 31.9438619 | 1.21314019 | 1.9815E-09 | 3.86E-07 |
| RnasE6 | 6.56501051 | 16.2070318 | 1.40848039 | 2.0498E-09 | 3.9385E-07 |
| Pi4k2b | 8.53958866 | 15.5458883 | 0.97382013 | 2.2604E-09 | 4.2843E-07 |
| Cth | 7.15494982 | 33.6687109 | 2.35881238 | 2.358E-09 | 4.4098E-07 |
| Trim35 | 19.016521 | 33.4483627 | 0.92676024 | 2.4585E-09 | 4.4784E-07 |
| Vim | 10.3259516 | 20.7216025 | 1.1177967 | 2.5649E-09 | 4.6123E-07 |
| Ighv1-75 | 211.47111 | 524.66666 | 1.42540097 | 2.6451E-09 | 4.6962E-07 |
| Pgghg | 3.03465101 | 6.91086231 | 1.29198015 | 2.7368E-09 | 4.7984E-07 |
| Igkv3-7 | 88.0774887 | 390.419959 | 2.23318595 | 3.0091E-09 | 5.2105E-07 |
| Fut1 | 3.32197449 | 11.546848 | 1.90485349 | 3.1433E-09 | 5.2677E-07 |
| Mvb12b | 1.98977647 | 5.57786571 | 1.60793752 | 3.1426E-09 | 5.2677E-07 |
| Igkv3-12 | 211.75589 | 469.149729 | 1.26373459 | 3.3043E-09 | 5.4524E-07 |
| Rab3d | 2.99415195 | 9.6165486 | 1.79447135 | 3.492E-09 | 5.6952E-07 |
| Ighv5-6 | 222.340625 | 498.126916 | 1.27894106 | 3.547E-09 | 5.7185E-07 |
| Dynlt3 | 13.0618782 | 23.145537 | 0.93705408 | 3.7139E-09 | 5.853E-07 |
| Ighv1-72 | 356.730241 | 683.831458 | 1.05372924 | 3.8906E-09 | 6.0632E-07 |
| MagEh1 | 2.21778289 | 10.4676177 | 2.36410193 | 3.9542E-09 | 6.0948E-07 |
| Gpt2 | 6.76469786 | 16.8743894 | 1.43288768 | 4.2146E-09 | 6.4254E-07 |
| Gpr107 | 3.26184279 | 5.67661715 | 0.91165444 | 4.4574E-09 | 6.7225E-07 |
| Psd3 | 0.90277054 | 2.11359823 | 1.3441571 | 4.5302E-09 | 6.7596E-07 |
| PdE4dip | 4.33027968 | 8.79605459 | 1.13419308 | 4.6921E-09 | 6.9172E-07 |
| Rap1gap2 | 2.37365066 | 7.12641108 | 1.68732496 | 4.8627E-09 | 6.9596E-07 |
| Ighv9-3 | 845.080869 | 1823.04019 | 1.22378646 | 5.1931E-09 | 7.3574E-07 |
| Sla | 10.3130531 | 33.0893294 | 1.7887961 | 5.4657E-09 | 7.6662E-07 |
| Arfgap1 | 13.6506179 | 23.2193189 | 0.88013341 | 5.9282E-09 | 7.922E-07 |
| Ighv7-1 | 57.8715547 | 185.210762 | 1.78376919 | 5.9305E-09 | 7.922E-07 |
| Atp1b1 | 2.2660117 | 9.54716438 | 2.17686473 | 5.9887E-09 | 7.9243E-07 |

| | | | | | |
|---|---|---|---|---|---|
| Wipi1 | 3.7306877 | 10.6721124 | 1.61168347 | 6.1487E-09 | 8.06E-07 |
| Ighv1-64 | 617.173425 | 1204.09356 | 1.07509134 | 6.2841E-09 | 8.1328E-07 |
| Igkv2-109 | 287.82215 | 538.623955 | 1.01780736 | 6.6414E-09 | 8.4685E-07 |
| Chst12 | 17.2410663 | 34.2471491 | 1.10523494 | 6.9979E-09 | 8.7636E-07 |
| Rcn1 | 5.95526686 | 10.9502489 | 0.98721985 | 6.9631E-09 | 8.7636E-07 |
| Cxcr4 | 34.2194442 | 71.7828985 | 1.17502586 | 7.275E-09 | 9.0301E-07 |
| TcEal9 | 105.81069 | 187.609369 | 0.94026669 | 7.3757E-09 | 9.0746E-07 |
| Tbc1d20 | 14.202912 | 22.7946966 | 0.79525867 | 7.8723E-09 | 9.5187E-07 |
| Yipf3 | 26.4038598 | 51.001768 | 1.06570332 | 8.1676E-09 | 9.7914E-07 |
| Ighv1-59 | 136.162105 | 282.814039 | 1.17170051 | 8.3655E-09 | 9.9436E-07 |
| Hspa2 | 6.14059259 | 13.8861053 | 1.2802731 | 8.9608E-09 | 1.0262E-06 |
| ManEa | 37.8668744 | 73.2119093 | 1.06531703 | 8.9992E-09 | 1.0262E-06 |
| SEc31a | 38.1036651 | 66.136888 | 0.90935517 | 9.1265E-09 | 1.0323E-06 |
| CrEb3 | 16.5128894 | 29.5790395 | 0.9496466 | 9.3113E-09 | 1.0365E-06 |
| Jdp2 | 1.19205083 | 6.32176305 | 2.49753902 | 9.2812E-09 | 1.0365E-06 |
| Coro2b | 2.28618196 | 5.05399866 | 1.25600966 | 9.5561E-09 | 1.0554E-06 |
| 4933421O10Rik | 2.28006423 | 5.15718667 | 1.28033209 | 9.8643E-09 | 1.0689E-06 |
| Yipf6 | 7.95407591 | 12.6897598 | 0.7875543 | 9.8369E-09 | 1.0689E-06 |
| Ighv4-1 | 99.3378797 | 302.735266 | 1.71218446 | 9.9876E-09 | 1.0694E-06 |
| Soat2 | 0.75885652 | 4.18559775 | 2.59352619 | 1.0314E-08 | 1.0959E-06 |
| Fam214a | 9.33662461 | 21.2478234 | 1.30747877 | 1.1086E-08 | 1.1433E-06 |
| Sar1a | 61.5542207 | 95.0026689 | 0.7386418 | 1.1042E-08 | 1.1433E-06 |
| Tlr4 | 2.18138918 | 4.80452863 | 1.2532204 | 1.1078E-08 | 1.1433E-06 |
| Ighv5-12 | 83.3384642 | 165.019472 | 1.09839051 | 1.1459E-08 | 1.1488E-06 |
| Raph1 | 0.36421368 | 1.1134717 | 1.70529336 | 1.1518E-08 | 1.1488E-06 |
| Sh3bp5l | 4.36236038 | 7.68444982 | 0.92495883 | 1.1222E-08 | 1.1488E-06 |
| Trp53inp1 | 49.4207807 | 157.939944 | 1.79681397 | 1.1549E-08 | 1.1488E-06 |
| Man1a | 63.5384161 | 108.541952 | 0.88776836 | 1.212E-08 | 1.1972E-06 |
| Dhrs7 | 9.05339474 | 23.5011877 | 1.48325556 | 1.2212E-08 | 1.1978E-06 |
| Asns | 67.3838875 | 158.609161 | 1.35761004 | 1.2423E-08 | 1.21E-06 |
| Calu | 89.7493436 | 142.866698 | 0.78412488 | 1.3481E-08 | 1.2951E-06 |
| Pofut2 | 3.77669546 | 6.54083503 | 0.90289031 | 1.3421E-08 | 1.2951E-06 |
| MEtrnl | 2.07898767 | 5.48153633 | 1.5115834 | 1.3847E-08 | 1.3212E-06 |
| Ighv6-6 | 494.366889 | 1029.03481 | 1.16762602 | 1.4143E-08 | 1.3313E-06 |
| Nbas | 22.376612 | 40.3065936 | 0.96697261 | 1.41E-08 | 1.3313E-06 |
| March8 | 1.14904799 | 2.59744859 | 1.27977824 | 1.4717E-08 | 1.367E-06 |
| Ttc39c | 1.04253478 | 3.39717741 | 1.79315649 | 1.5201E-08 | 1.3935E-06 |
| Golim4 | 3.32533095 | 5.96200264 | 0.95450019 | 1.5589E-08 | 1.4107E-06 |
| Rhbdd1 | 4.06929643 | 8.09558848 | 1.10001075 | 1.5854E-08 | 1.4255E-06 |
| Ighv1-20 | 31.0580896 | 86.6437394 | 1.60413591 | 1.6336E-08 | 1.4493E-06 |
| Igkv1-122 | 105.305975 | 274.041408 | 1.48509659 | 1.6516E-08 | 1.4493E-06 |
| Igkv8-30 | 806.690665 | 2738.69763 | 1.85815638 | 1.6636E-08 | 1.4493E-06 |
| Il5ra | 20.7553655 | 39.0275117 | 1.01678188 | 1.6576E-08 | 1.4493E-06 |
| Mospd1 | 10.6793553 | 21.7265592 | 1.12801229 | 1.6431E-08 | 1.4493E-06 |
| Ppp1r15a | 15.8581232 | 27.9014134 | 0.93147207 | 1.6837E-08 | 1.4578E-06 |

| | | | | | |
|---|---|---|---|---|---|
| Acbd3 | 11.6325609 | 18.0465504 | 0.74797655 | 1.7142E-08 | 1.4661E-06 |
| Gm14137 | 0.90074637 | 4.01583048 | 2.25172185 | 1.7088E-08 | 1.4661E-06 |
| Gm30211 | 12.2902208 | 24.1378561 | 1.08916142 | 1.7861E-08 | 1.5092E-06 |
| Ctla4 | 13.9760963 | 38.1724462 | 1.57311333 | 1.8213E-08 | 1.5297E-06 |
| IpcEf1 | 1.64955262 | 4.63477949 | 1.58437819 | 1.8441E-08 | 1.5305E-06 |
| Slc7a3 | 17.9017152 | 43.2030852 | 1.38682823 | 1.9278E-08 | 1.5906E-06 |
| Sqstm1 | 95.6690791 | 233.071757 | 1.39260349 | 1.9392E-08 | 1.5906E-06 |
| Rgs12 | 0.34547675 | 0.92013488 | 1.52553039 | 1.9682E-08 | 1.5986E-06 |
| TmEm167b | 11.8839628 | 22.8986086 | 1.06340183 | 1.9717E-08 | 1.5986E-06 |
| Igkv1-88 | 106.94773 | 307.237129 | 1.63506914 | 2.0161E-08 | 1.6159E-06 |
| Ighv2-5 | 56.3886886 | 145.854396 | 1.47428516 | 2.0386E-08 | 1.6233E-06 |
| Slc12a4 | 6.60095176 | 11.5270568 | 0.91251442 | 2.0485E-08 | 1.6233E-06 |
| ErlEc1 | 24.5122791 | 45.4963191 | 1.00650571 | 2.0912E-08 | 1.6246E-06 |
| Gadd45a | 10.2008496 | 28.4897161 | 1.58648332 | 2.0965E-08 | 1.6246E-06 |
| Tlr2 | 3.61433228 | 7.75343662 | 1.20865949 | 2.069E-08 | 1.6246E-06 |
| Hmox1 | 13.6122844 | 24.6654684 | 0.9677937 | 2.1635E-08 | 1.6673E-06 |
| Ddit3 | 6.14854641 | 17.8754547 | 1.62865743 | 2.2384E-08 | 1.7047E-06 |
| Igkv1-133 | 81.010718 | 234.867302 | 1.65210838 | 2.2484E-08 | 1.7047E-06 |
| Slc50a1 | 22.2875933 | 46.7446363 | 1.17191099 | 2.2465E-08 | 1.7047E-06 |
| Slc26a2 | 3.47573678 | 5.813726 | 0.85498431 | 2.2926E-08 | 1.7288E-06 |
| HErpud1 | 197.528765 | 357.204429 | 0.96986954 | 2.3247E-08 | 1.7402E-06 |
| Ighv14-3 | 271.711793 | 695.678222 | 1.47525945 | 2.3449E-08 | 1.7402E-06 |
| REEp6 | 0.46224492 | 2.50082952 | 2.52070194 | 2.3728E-08 | 1.7424E-06 |
| Igkv5-45 | 78.6227622 | 221.9713 | 1.58989651 | 2.4123E-08 | 1.7614E-06 |
| Unc13d | 6.76383252 | 12.2349884 | 0.96284153 | 2.4237E-08 | 1.7614E-06 |
| Ighv13-2 | 87.1523552 | 238.350201 | 1.54552775 | 2.5359E-08 | 1.8162E-06 |
| St3gal1 | 13.8681071 | 22.7931321 | 0.83204387 | 2.538E-08 | 1.8162E-06 |
| EthE1 | 10.5008401 | 19.2597169 | 0.98359719 | 2.6478E-08 | 1.8852E-06 |
| Gfpt1 | 26.1110851 | 44.5335744 | 0.88472744 | 2.7662E-08 | 1.9399E-06 |
| Inpp4b | 2.12924116 | 4.45148634 | 1.18314577 | 2.7625E-08 | 1.9399E-06 |
| REln | 0.26533184 | 0.77656231 | 1.6720055 | 2.7612E-08 | 1.9399E-06 |
| Il2ra | 5.12527102 | 10.4770576 | 1.13580663 | 2.864E-08 | 1.9887E-06 |
| Glipr2 | 27.8629379 | 50.970921 | 0.98469683 | 2.882E-08 | 1.9913E-06 |
| Gpr180 | 15.8570288 | 27.2937074 | 0.89726059 | 2.9468E-08 | 2.0261E-06 |
| Jchain | 714.95209 | 1411.49132 | 1.09028911 | 2.998E-08 | 2.0413E-06 |
| Ighv1-76 | 317.430465 | 612.753878 | 1.06069791 | 3.1154E-08 | 2.1008E-06 |
| Snn | 8.43828778 | 20.35443 | 1.3834221 | 3.1645E-08 | 2.1136E-06 |
| Kcnk6 | 9.75839385 | 18.5056735 | 1.03935722 | 3.1996E-08 | 2.1168E-06 |
| Cdkn1a | 1.84936633 | 5.41871871 | 1.65105609 | 3.2588E-08 | 2.1359E-06 |
| Slc7a7 | 5.86009167 | 13.7831684 | 1.35063664 | 3.2529E-08 | 2.1359E-06 |
| Tvp23b | 19.3302306 | 32.6427742 | 0.86499699 | 3.3936E-08 | 2.1834E-06 |
| Gpx3 | 6.16358458 | 30.3180393 | 2.38096243 | 3.4574E-08 | 2.1855E-06 |
| Trib2 | 8.63961301 | 15.8867067 | 0.99702627 | 3.4592E-08 | 2.1855E-06 |
| GnE | 14.0232896 | 27.4199704 | 1.08280767 | 3.4933E-08 | 2.1938E-06 |
| PtprE | 1.28405611 | 2.41783789 | 1.02138742 | 3.5308E-08 | 2.1996E-06 |

| | | | | | |
|---|---|---|---|---|---|
| Bsdc1 | 6.21022664 | 10.3785412 | 0.85146346 | 3.6098E-08 | 2.2227E-06 |
| Igkv15-103 | 400.681408 | 712.142057 | 0.93951616 | 3.6132E-08 | 2.2227E-06 |
| Ighv8-12 | 303.281523 | 740.773658 | 1.39210929 | 3.683E-08 | 2.2558E-06 |
| NdEl1 | 15.4372666 | 24.3417372 | 0.76852059 | 3.7821E-08 | 2.2866E-06 |
| Inpp1 | 5.99734133 | 11.2262362 | 1.01543328 | 3.9591E-08 | 2.363E-06 |
| Crlf2 | 16.2972407 | 38.3597722 | 1.33544547 | 3.9815E-08 | 2.3663E-06 |
| Lipc | 1.32408214 | 4.41486074 | 1.84464392 | 4.0313E-08 | 2.3858E-06 |
| Mon2 | 9.23027003 | 15.2361839 | 0.83883763 | 4.0637E-08 | 2.3909E-06 |
| Ttc39b | 3.26640869 | 5.19248685 | 0.78041416 | 4.1332E-08 | 2.4155E-06 |
| Cul7 | 1.00084922 | 3.35346293 | 1.83269295 | 4.2245E-08 | 2.4479E-06 |
| Igkv4-74 | 48.8608106 | 119.216005 | 1.3949953 | 4.2169E-08 | 2.4479E-06 |
| Mib2 | 5.1346544 | 14.1981036 | 1.58319683 | 4.2409E-08 | 2.4479E-06 |
| Cog6 | 15.9612547 | 24.0849671 | 0.70686805 | 4.3689E-08 | 2.4985E-06 |
| Ighv9-2 | 28.7142745 | 92.7714972 | 1.80983639 | 4.3613E-08 | 2.4985E-06 |
| Trib3 | 5.85419084 | 21.1884096 | 1.96250387 | 4.4533E-08 | 2.4985E-06 |
| Tspan13 | 77.8001979 | 131.917964 | 0.87743391 | 4.451E-08 | 2.4985E-06 |
| SEl1l | 74.0349733 | 141.875679 | 1.05635002 | 4.5223E-08 | 2.5171E-06 |
| Slc1a4 | 7.04781183 | 11.4766503 | 0.81518158 | 4.701E-08 | 2.5959E-06 |
| Atp9a | 1.57194625 | 3.27298766 | 1.16168884 | 4.8779E-08 | 2.6212E-06 |
| Pdxdc1 | 12.350447 | 19.4733908 | 0.77042941 | 4.8824E-08 | 2.6212E-06 |
| SErpina3g | 3.58522835 | 8.78025916 | 1.4277383 | 4.8962E-08 | 2.6212E-06 |
| Slc35a2 | 7.31890771 | 11.5176053 | 0.7670044 | 4.8527E-08 | 2.6212E-06 |
| Susd6 | 5.67195617 | 10.4634998 | 0.99234617 | 4.8388E-08 | 2.6212E-06 |
| TmEm123 | 135.814148 | 242.749425 | 0.95188896 | 4.874E-08 | 2.6212E-06 |
| NEo1 | 0.07054629 | 0.62612851 | 3.20581061 | 5.0577E-08 | 2.677E-06 |
| Gabarap | 91.102466 | 166.819847 | 0.98241311 | 5.1323E-08 | 2.7062E-06 |
| Rab6a | 29.6031566 | 43.8498005 | 0.67849197 | 5.3167E-08 | 2.793E-06 |
| Prrc1 | 12.8510305 | 20.6633024 | 0.80155072 | 5.4544E-08 | 2.8278E-06 |
| Rnf103 | 3.49859977 | 5.96573581 | 0.88511906 | 5.4837E-08 | 2.8278E-06 |
| RnpEpl1 | 4.81889328 | 7.5543354 | 0.76074068 | 5.4383E-08 | 2.8278E-06 |
| Endod1 | 6.15316971 | 10.1126632 | 0.83226657 | 5.5836E-08 | 2.8582E-06 |
| Praf2 | 5.75380578 | 10.8659631 | 1.02993185 | 5.6876E-08 | 2.8746E-06 |
| Spint2 | 7.34653617 | 12.2139734 | 0.84493543 | 5.7105E-08 | 2.8746E-06 |
| Yif1a | 21.6809215 | 38.5979021 | 0.94026825 | 5.718E-08 | 2.8746E-06 |
| Lmo4 | 15.1512899 | 24.5464423 | 0.80920763 | 5.7623E-08 | 2.8865E-06 |
| Dnajc1 | 3.60139592 | 5.54067935 | 0.7348424 | 6.2023E-08 | 3.0417E-06 |
| Itpripl2 | 0.12803576 | 0.87919737 | 2.91518289 | 6.247E-08 | 3.0424E-06 |
| Slc39a13 | 4.27827583 | 7.45111603 | 0.91148684 | 6.3512E-08 | 3.0718E-06 |
| Pja2 | 7.85630026 | 13.6083297 | 0.90560669 | 6.4489E-08 | 3.1083E-06 |
| Gm16350 | 23.5919258 | 55.308813 | 1.33667862 | 6.6362E-08 | 3.1745E-06 |
| Prxl2c | 6.00134439 | 10.7218524 | 0.94727964 | 6.6768E-08 | 3.1745E-06 |
| Ighv5-15 | 25.9141269 | 93.8358001 | 1.95951179 | 6.9233E-08 | 3.2696E-06 |
| Igkv8-21 | 253.683194 | 562.517456 | 1.25460324 | 7.0511E-08 | 3.3107E-06 |
| Il3ra | 2.82948133 | 7.56597796 | 1.52696051 | 7.0576E-08 | 3.3107E-06 |
| Mthfr | 5.233514 | 8.95630871 | 0.88638671 | 7.1059E-08 | 3.3112E-06 |

| | | | | | |
|---|---|---|---|---|---|
| Igkv4-86 | 87.0225436 | 199.965094 | 1.30964676 | 7.1343E-08 | 3.3135E-06 |
| Parp4 | 7.50761629 | 13.5623984 | 0.96836996 | 7.2066E-08 | 3.325E-06 |
| CEbpg | 12.43347 | 21.0710695 | 0.87044679 | 7.541E-08 | 3.4341E-06 |
| Bckdha | 9.20843982 | 20.0079007 | 1.22548147 | 7.6162E-08 | 3.4571E-06 |
| DErl1 | 129.802242 | 211.170482 | 0.81522259 | 7.7354E-08 | 3.4774E-06 |
| AU040320 | 8.82820863 | 15.8680335 | 0.96071512 | 7.7704E-08 | 3.4814E-06 |
| Bmf | 5.67799777 | 13.2170348 | 1.32586415 | 7.7938E-08 | 3.4814E-06 |
| NfE2l1 | 34.552628 | 52.8959632 | 0.7293996 | 8.3007E-08 | 3.6612E-06 |
| Cln8 | 1.85300726 | 3.56768818 | 1.05099426 | 8.3783E-08 | 3.6838E-06 |
| HEatr5a | 1.83001089 | 3.09829889 | 0.86998659 | 8.7052E-08 | 3.7396E-06 |
| Mcrip1 | 21.7015057 | 41.1268732 | 1.0288351 | 8.7224E-08 | 3.7396E-06 |
| Mknk1 | 15.440431 | 30.7643332 | 1.09712445 | 8.6447E-08 | 3.7396E-06 |
| SErf2 | 14.1600435 | 23.0294803 | 0.8141757 | 8.7451E-08 | 3.7396E-06 |
| Zfp260 | 8.1133273 | 15.370286 | 1.04214298 | 8.68E-08 | 3.7396E-06 |
| SEcisbp2l | 6.25666729 | 10.5971096 | 0.87534107 | 8.8421E-08 | 3.7696E-06 |
| Aig1 | 0.94331833 | 2.25313561 | 1.35118953 | 8.8851E-08 | 3.7717E-06 |
| Igkv5-43 | 770.267051 | 1568.91977 | 1.13458543 | 8.9395E-08 | 3.7717E-06 |
| H6pd | 6.36635909 | 10.2183714 | 0.79372077 | 9.0285E-08 | 3.7914E-06 |
| Golga2 | 16.4318795 | 27.286247 | 0.84866278 | 9.2083E-08 | 3.8554E-06 |
| Igkv4-58 | 88.4913289 | 188.755464 | 1.19631903 | 9.313E-08 | 3.8876E-06 |
| Gpr155 | 2.52403384 | 4.25907048 | 0.86477407 | 9.7929E-08 | 4.0398E-06 |
| Trib1 | 11.316922 | 21.1980173 | 1.02325024 | 9.9125E-08 | 4.0772E-06 |
| Srgn | 141.106116 | 248.672381 | 0.92835613 | 1.0119E-07 | 4.1306E-06 |
| Yipf4 | 17.9864186 | 26.87745 | 0.69029546 | 1.0217E-07 | 4.1306E-06 |
| Zfp324 | 4.62140848 | 7.11153232 | 0.73296432 | 1.0151E-07 | 4.1306E-06 |
| BEt1l | 4.86459743 | 8.43023078 | 0.89663215 | 1.0422E-07 | 4.1723E-06 |
| Prkcz | 0.73956701 | 1.61513008 | 1.23758219 | 1.0458E-07 | 4.1723E-06 |

**Supplementary Table 3. List of antibodies used in flow cytometry, western blot and immunofluorescence**

| Antibody or Dye | Clone | Host/Isotype | Supplier |
|---|---|---|---|
| **Flow cytometry** | | | |
| Anti-CD138 | 281-2 | Rat IgG2a, κ | BD Biosciences |
| Anti-CD45R/B220 | RA3-6B2 | Rat / IgG2a, kappa | BD Biosciences |
| Anti-CD19 | 1D3 | Rat IgG2a, κ | BD Biosciences |
| Anti-CD21/35 | 7G6 | Rat gG2b, κ | BD Biosciences |
| Anti-CD23 | B3B4 | Rat IgG2a, κ | BD Biosciences |
| Anti-CD93 | AA4.1 | Rat IgG2a, κ | BD Biosciences |
| Anti-IgM | II/41 | Rat IgG2a, κ | eBioscience |
| Anti-Taci | 8F10-3 | Rat IgG2a, κ | eBioscience |
| PE Annexin-V Apoptosis Detection Kit I | | | BD Pharmingen |
| V450-anti-cleaved caspase-3 | | | BD Biosciences |
| Anti-KI67 Alexa fluor 700 | B56 | mouse IgG1 | BD Biosciences |
| Dapi | | | |
| ERT | | | Invitrogen |
| lysoT | | | Invitrogen |
| mitoT green | | | Invitrogen |
| mitoT orange | | | Invitrogen |
| Golgi painter | | | abcam |
| Viability dye e506 | | | Invitrogen |
| Aqua zombie | | | Biolegend |
| **Immunofluorescence** | | | |
| Anti-Calnexin | | Rabbit IgG | Abcam |
| Anti-cytochrome c | | Rabbit IgG | Thermofisher |
| Anti-GM130 | | Rabbit IgG | Thermofisher |
| anti-Rabbit IgG Alexa fluor 555 | | Goat IgG | Thermofisher |
| anti-mouse IgM Alexa fluor 647 | | Goat IgG | Thermofisher |
| Hoescht33342 | | | Thermofisher |
| **Western blot** | | | |
| Anti-sec22b | | Rabbit IgG | Synaptic Systems |
| Anti-b actin | | Rabbit IgG | Cell signaling |
| Anti-Xbp1 | | Rabbit IgG | Cell Signaling |
| Anti-IgM | | Goat IgG | Southern Biotech |
| Anti-CD138 | | Rabbit IgG | Poteintech |
| Anti-Scfd1/rsly1 | | Rabbit IgG | Dr Jesse Hay, PMID: 14565970 |
| Anti-Uso1/p115 | | Rabbit IgG | Dr Jesse Hay, PMID: 25406594 |

| | | | |
|---|---|---|---|
| Anti-Stx5 | | Rabbit IgG | Synaptic Systems |
| Anti-Ykt6 | | Rabbit IgG | Dr Jesse Hay; PMID: 12589064 |
| Anti- Rabbit HRP | | Goat IgG | Jackson immuno research |
| **ELISA** | | | |
| Anti mouse Ig kappa HRP | | Goat IgG | Southern Biotech |
| Anti mouse IgM HRP | | Goat IgG | Southern Biotech |
| Anti mouse IgG1 HRP | | Goat IgG | Southern Biotech |
| Anti mouse IgA HRP | | Goat IgG | Southern Biotech |
| Anti mouse IgG3 HRP | | Goat IgG | Southern Biotech |

**Supplementary Table 4: List of primers used for Biomark based qPCR**

| | Gene | Reference/ Primers |
|---|---|---|
| Taqman assays | *Gapdh* | Mm99999915_g1 |
| | *Actb* | Mm01205647_g1 |
| | *Prdm1* | Mm00476128_m1 |
| | *Xbp1* | Mm00457357_m1 |
| | *Irf4* | Mm00516431_m1 |
| | *Pax5* | Mm00435501_m1 |
| | *Cd3e* | Mm00599684_g1 |
| | *Bach2* | Mm00464379_m1 |
| | *Cd93* | Mm00440239_g1 |
| | *Hspa5* | Mm00517691_m1 |
| | *Edem2* | Mm00467468_m1 |
| | *Dnajb9* | Mm01622956_s1 |
| | *Atg5* | Mm01187303_m1 |
| | *Traf2* | Mm00801978_m1 |
| | *Atg12* | Mm00503201_m1 |
| | *Atg16L1* | Mm00513085_m1 |
| | *Ddit3* | Mm01135937_g1 |
| | *Becn1* | Mm01265461_m1 |
| | *Herpud1* | Mm01249592_m1 |
| | *Edem1* | Mm00551797_m1 |
| | *Tnfrsf13b* | Mm03047441_m1 |
